# Using Extracellular miRNA Signatures to Identify Patients with LRRK2-Related Parkinson’s Disease

**DOI:** 10.1101/2023.11.06.565815

**Authors:** Luca Jannik Braunger, Felix Knab, Thomas Gasser

## Abstract

**Background:** Mutations in the Leucine Rich Repeat Kinase 2 gene are highly relevant in both sporadic and familial cases of Parkinson’s disease. Specific therapies are entering clinical trials but patient stratification remains challenging. Dysregulated microRNA expression levels have been proposed as biomarker candidates in sporadic Parkinson’s disease.

**Objective:** In this proof-of concept study we evaluate the potential of extracellular miRNA signatures to identify LRRK2-driven molecular patterns in Parkinson’s disease.

**Methods:** We measured expression levels of 91 miRNAs via RT-qPCR in ten individuals with sporadic Parkinson’s disease, ten *LRRK2* mutation carriers and eleven healthy controls using both plasma and cerebrospinal fluid. We compared miRNA signatures using heatmaps and t-tests. Next, we applied group sorting algorithms and tested sensitivity and specificity of their group predictions.

**Results:** miR-29c-3p was differentially expressed between *LRRK2* mutation carriers and sporadic cases, with miR-425-5p trending towards significance. Individuals clustered in principal component analysis along mutation status. Group affiliation was predicted with high accuracy in the prediction models (sensitivity up to 89%, specificity up to 70%). miRs-128-3p, 29c-3p, 223-3p and 424-5p were identified as promising discriminators among all analyses.

**Conclusions:** LRRK2 mutation status impacts the extracellular miRNA signature measured in plasma and separates mutation carriers from sporadic Parkinson’s disease patients. Monitoring LRRK2 miRNA signatures could be an interesting approach to test drug efficacy of LRRK2-targeting therapies. In light of small sample size, the suggested approach needs to be validated in larger cohorts.

## Introduction

The majority of familial PD (fPD) cases is caused by mutations in the Leucine Rich Repeat Kinase 2 (*LRRK2*) gene [1,2], with the G2019S mutation being the most common [3]. Despite its apparent role in PD, LRRK2 expression levels in the CNS are relatively low compared to peripheral tissues such as the lung and the kidneys, as well as blood monocytes [4,5]. Increased kinase activity is thought to be the cause of the disturbed cellular homeostasis observable in cell models of PD [6]. Most importantly, LRRK2 was shown to play a role in sporadic cases, with non-coding variants in the *LRRK2* gene increasing the risk of PD [7]. Further, sporadic PD (sPD) and fPD cases show overlapping clinical features [8]. Studying fPD cases therefore offers the opportunity to better understand the molecular pathogenesis of both fPD and sPD cases. Targeting multiple molecular pathways in selected PD patients with an individually formulated combination of drugs could be a potential strategy for disease modification [9]. However, this strategy necessitates not only the discovery of particular druggable pathways, but also the careful selection of patients who are most likely to benefit from a given treatment, ideally at an early or even asymptomatic stage in the course of their disease. While aberrant LRRK2 activity represents such a molecular target and LRRK2 inhibitors are approaching clinical testing [10], [11], pre-selecting patients as well as monitoring target engagement and drug efficacy remain challenging. There are no established biomarkers available that can aid in the early detection of disease onset before significant neurodegeneration occurs.

Recently, microRNAs (miRNAs), small non-coding RNA molecules that are involved in the posttranscriptional regulation of gene expression [12], have been intensively studied as potential biomarkers for neurodegenerative diseases, including PD [13,14]. So far, most biomarker studies using miRNAs have focused on the differentiation of sPD patients from healthy controls (HC), while in the present proof-of-concept study we aimed to test how well extracellular miRNAs can be used to reliably differentiate between sPD patients and *LRRK2* mutation carriers (LRRK2_MC_). Since the distinction between fPD and sPD is possible by using more stringent methods, such as sequencing, in the long run we aim to identify individuals with relevant involvement of LRRK2-dependent pathways among the sPD population. However, in a first step we decided to define what constitutes a molecular LRRK2 fingerprint by comparing the extracellular miRNA signatures of fPD and sPD cases. Given that LRRK2 was described to change the cellular miRNAome [15], we hypothesized that mutations in the *LRRK2* gene also alter the extracellular miRNA signature in LRRK2_MC_.

## Materials and Methods

### Experimental Design

In a first step, we quantified the expression levels of 91 extracellular miRNAs from a cohort of ten sPD, ten fPD and eleven HC in both plasma and CSF using RT-qPCR (Fig. 1). After processing of raw Ct values and calculation of log2 fold change values (log2fc), as described in more detail in the respective method section, we performed t-tests to identify single miRNAs with the potential of discriminating between groups. We further applied group sorting algorithms such as principal component analysis (PCA), least absolute shrinkage and selection operator (LASSO) and random forest (RF) using the calibrated Ct values of all miRNAs that could be reliably detected throughout the cohort. The ability to differentiate between sPD patients and LRRK2_MC_ with and without PD was assessed for each sorting algorithm by evaluating their sensitivity and specificity. Finally, we selected those miRNAs that had a significant impact on the performance of each model.

**Figure 1:**
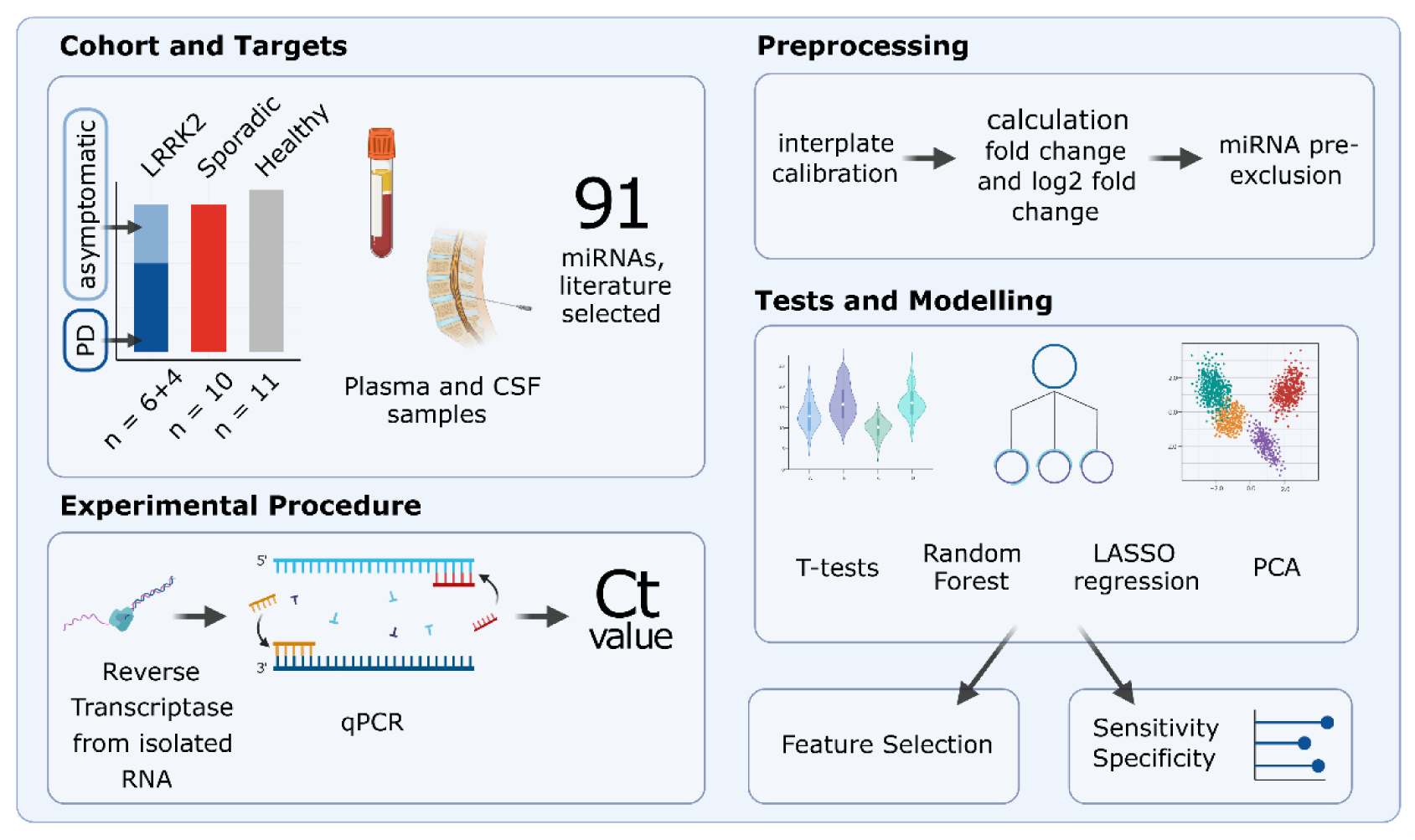
Study Design Overview. The experimental cohort compromised 31 patients categorized into three groups: individuals carrying a *LRRK2* mutation, sporadic PD patients and healthy controls. A total of 91 miRNAs, selected by reviewing relevant literature, were quantified in CSF and plasma samples using reverse transcription and qPCR. Subsequently, the obtained raw Ct values were calibrated and fc and log2fc values were calculated. miRNAs that were expressed only in subgroups were excluded. Group differences were then analyzed using t-tests, PCA, Random Forest and LASSO regression. Created with BioRender.com.

### Cohort Design

For each LRRK2_MC_, one sPD and one HC was added to the study cohort. To obtain homogenous groups, individuals were selected based on age, gender, and disease duration. As the histopathological complexity of PD likely increases over time and patients with long disease durations might no longer display pathologies specific to mutation status rather than showing features common to all PD patients, disease durations were kept as short as possible. The group of LRRK2_MC_ initially included a PD patient carrying two *LRRK2* variants (N1437S, S1647). Given that these polymorphisms were not reported to be pathogenic, we finally decided to exclude this patient from the analysis. The final cohort therefore comprised three groups (Table 1): Ten LRRK2_MC_ (fPD patients: n=6 (*LRRK2* G2019S: n = 3; *LRRK2* G2019S/G1819: n=1, *LRRK2* R1441C: n=1; *LRRK2* I2020T: n=1), asymptomatic G2019S mutation carriers: n=4), ten sPD patients and eleven HCs. After performing PCA as described in more detail in the respective method section, the data set of one of the asymptomatic LRRK2_MC_ appeared as a technical or biological outlier and was removed from the plasma dataset before further analysis (Fig. S1). Clinical features are reported in Table 1. The researcher performing the experiments was blinded to the group annotation.

**Table 1:** Overview of clinical data.

| Disease status | *LRRK2* mutated |  | Sporadic PD (n = 10) | Healthy Control (n = 11) |
| --- | --- | --- | --- | --- |
|  | PD (n = 6) | Asymptomatic (n = 4) |  |  |
| **Age (years)** | 66.8 ± 8.7 | 52.3 ± 19.6 | 64.5 ± 10.9 | 60.1 ± 14.5 |
| **Gender (m:f)** | 1:5 | 0:4 | 2:8 | 2:9 |
| **DD (years)** | 8.8 ± 3.6 | - | 7 ± 4.1 | - |
| **MoCA** | 25 ± 4.1 (5/6) | 28.5 ± 1.5 | 25.3 ± 3.7 (9/10) | 28.3 ± 1.5 (4/11) |
| **UPDRS-III** | 21.7 ± 8.2 | 1.0 ± 0.8 (3/4) | 17.4 ± 10.1 (8/10) | 0.7 ± 0.9 (3/11) |
| **LEDD (mg)** | 612.8 ± 295.6 | 0.0 ± 0.0 | 357.5 ± 380.0 | 0.0 ± 0.0 |

DD: Disease duration, MoCA: Montreal-Cognitive-Assessment, UPDRS-III: Unified Parkinson's Disease Rating Scale, LEDD: L-Dopa equivalent daily dose. Age, Disease duration, MoCA, UPDRS-III and LEDD stated as mean value with standard deviation. MoCA and UPDRS-III were not accessible for some patients and numbers in brackets give fraction of accessible data.

### miRNA selection and qPCR panel design

We scanned the literature for miRNAs that were previously reported to be dysregulated in the context of *LRRK2* mutations, functionally relevant to LRRK2 or generally dysregulated in the context of PD. Next, we selected a total of 91 miRNAs to be included in our customized qPCR panel (Table S1). Selection was based on a set of criteria, including the number of relevant studies reporting dysregulation and a preference for miRNAs assessed in CSF and plasma samples over miRNAs reported in brain tissue or animal models. Additionally, miRNAs associated with LRRK2-linked PD were given priority over those associated with sporadic PD or other neurodegenerative diseases. For quality control purposes, we included a set of spike-in controls. Those included UniSpike 2 and 4 from the QIAGEN RNA Spike-In Kit (Cat. No.: 339390, QIAGEN, Hilden, Germany), which were used to monitor RNA isolation efficacy, UniSpike 6 (included in the QIAGEN miRCURY LNA RT Kit) to verify the efficacy of reverse transcription (RT) and UniSpike 3 (pre-applied primers on the custom qPCR plates) for inter-plate calibration. Additionally, a blank spot containing no primer was included and functioned as a negative control.

### Collection and Storage of Biofluids from Study Participants

Blood and CSF samples were collected from individuals according to standardized protocols. Within 90 minutes after collection from the individual, the collected material was brought from the medical facilities to the Neuro-Biobank Tuebingen and immediately processed. CSF and EDTA plasma were centrifuged at 2000g for 10 minutes. Samples were then transferred to a 15 ml Falcon tube, mixed thoroughly by vortexing, and aliquoted to cryotubes. Finally, both CSF and plasma cryotubes were frozen and stored at -80°C in the biobank. Before RNA isolation, samples were slowly thawed on ice.

### RNA isolation

Extracellular RNA was isolated from plasma and CSF using the QIAGEN miRNeasy Serum/Plasma Advanced Kit (Cat. No: 217204, QIAGEN) and following the vendor’s instructions with slight modifications. Briefly, 200 µl of plasma or 400 µl of CSF were used as a starting volume and subsequent reaction volumes were adjusted accordingly. After addition of RPL buffer and incubation for three minutes, we added 1 µl of the spike-in mix provided in the QIAGEN RNA Spike-In Kit (Cat. No.: 339390, QIAGEN) to the tube, in order to monitor the efficacy of the RNA isolation. Finally, RNA elution volume was reduced to 7 µl of RNase-free water.

### Reverse Transcription

The RT reaction was performed using the QIAGEN miRCURY LNA RT Kit (Cat. No.: 339340, QIAGEN). The reaction mix included 2 µl of 5x miRCURY SYBR® Green RT Reaction Buffer, 1 µl of 10x miRCURY RT Enzyme Mix, 4.5 µl of RNase-free water, 0.5 µl of UniSpike 6 RNA spike-in and 2 µl of template RNA per sample. After brief centrifugation, tubes were incubated at 42 °C for 60 minutes, followed by 5 minutes of incubation at 95 °C, which inactivated the RT reaction. Finally, samples were diluted 1:40 using RNase-free water.

### qPCR

qPCR was performed using QIAGEN miRCURY LNA miRNA Custom PCR Panels (Catalog No.: 339330, QIAGEN) along with the QIAGEN miRCURY LNA SYBR Green PCR Kit (Cat. No.: 339347, QIAGEN). The reaction mix included 515 µl of 2x miRCURY SYBR® Green Master Mix, 115 µl of RNase-free water and 400 µl of the cDNA template per sample. After pipetting 10 µl into each reaction-well, which already contained the dried primers, the plates were sealed and vortexed for one minute. After brief centrifugation, plates were incubated for ten minutes at room temperature to allow primers to resolve. Finally, plates were vortexed for one minute and briefly centrifuged. Amplification and quantification were performed using a LightCycler® 480 instrument (Roche, Basel, Switzerland). The cycling program included a two-minute heat activation step at 95 °C, followed by 45 cycles of denaturation at 95 °C for ten seconds and annealing at 56 °C for 60 seconds. To ensure specificity of amplifications, melting points of the amplified products were analyzed (Table S2 and S3).

### Processing of raw data

Before applying statistical analysis, raw data were processed and Ct and fold-change values (fc) were calculated (Fig. S2): First, raw data from the qPCR were converted using the LC480Conversion software to make them readable for LinRegPCR (Version 11.0, [16]), which was used for calculating Ct values. Further processing and analysis of data was performed in R (Version 4.2.2) using RStudio (Version 2023.03.0+386) [17]. To account for variations between plates, the Ct values were normalized using an interplate calibration factor. The calibration factor for each plate was calculated by subtracting the mean Ct value of UniSpike 3 reactions from all plates from the mean Ct value of all UniSpike 3 reactions in a given plate. All Ct values of the respective plate were then calibrated by subtracting the respective calibration factor from all Ct values in that plate. After calibration, Ct values > 40 were considered unspecific and excluded from the dataset. If a miRNA was not specifically amplified in more than one individual, it was removed completely from the analysis, after confirming that missing values occurred equally over all groups (Fig. S3). This resulted in 58 remaining miRNAs for the plasma dataset and eleven miRNAs for the CSF dataset.

Next, the fc and log2fc values were calculated using the 2^-ΔΔCt^ method (Fig. S4) [18]. In a first step, for each individual we independently calculated the mean Ct by adding all Ct values of every quantified miRNA and dividing it by the number of quantified miRNAs, which resulted in a *mean_Ct_* for each individual. We then computed ΔCt for each miRNA and individual by subtracting the *mean_Ct_* of the respective individual from every miRNA quantified in that individual. This resulted in multiple *ΔCt_miR-n_* per individual, where *n* represents the specific miRNA. The entire group of healthy controls was used as a reference group. Subsequently, for each miRNA we calculated the mean of *ΔCt_miR-n_*, only using *ΔCt_miR-n_* from the HCs (*mean ΔCt_miR-n-healthy_*). Next, ΔΔCt for each miRNA and individual was determined by subtracting the *mean ΔCt_miR-n-healthy_* from the patient’s *ΔCt_miR-n_*. Finally, the fold change was obtained using the following formula: fold change = 2^-ΔΔCt^. To achieve a linear scale with symmetry around zero, log2fc was used for the heatmaps and t-tests. Data from the HCs were used only as a reference for calculating the fold change values. Since the focus of this study was put on the identification of a LRRK2-driven molecular patterns and the discrimination between fPD and sPD cases, in the subsequent analyses, data from the HCs were not analyzed further.

### Data Visualization

Heatmaps visualizing either the log2fc or calibrated Ct values were created using the *pheatmap* package in R [19]. Log2fc and Ct values were scaled row-wise to z-scores using the *scale* function while ignoring missing values. After confirming normality of the data via assessing QQ-Plots, an unpaired t-test using the log2fc values was performed for each miRNA, comparing the sPD group with the LRRK2_MC_. The p-value threshold to determine significance was corrected for multiple testing by dividing by the number of tests. The distribution of p-values was examined and visualized via histograms. For miRNAs that the t-test revealed to be significantly differentially expressed between the groups, ROC analysis was performed. PCA was performed on calibrated Ct values using the built-in *prcomp* function of R. As the function does not accept missing values, they were imputed by setting them to the mean Ct value from all individuals. Two-dimensional plots were generated by graphing PC1 and PC2 values.

### Least Absolute Shrinkage and Selection Operator regression models

The *caret* package [20] was used for building the LASSO regression models. For missing values, the mean of the Ct values of all patients was used. We selected sensitivity and specificity in differentiating between the sPD and LRRK2_MC_ group, determined by Leave-One-Out-Cross-Validation (LOOCV) [21], as our experimental read-outs.

### Random forest Prediction Model

For building a RF model that predicts group affiliation to either the sPD or the LRRK2_MC_ group, the *randomForest* package [22] was used. To obtain mean values of sensitivity and specificity along with 95% confidence intervals, 100 models were constructed with each model containing 200 trees. For the RF model we did not use LOOCV, as sensitivity and specificity of the classification is already tested on out-of-bag samples reducing the risk of over-fitting. Along with this, we generated proximity heatmaps describing similarity of patient pairs based on the number of trees that classify those two patients in the same terminal node. Missing values were computed using the *rfImpute* function. The Gini coefficient’s mean decrease, which describes each decision node in terms of classification accuracy, was used to evaluate the importance of each variable. High purity of a node results in a low Gini coefficient. Consequently, variables that are more crucial for distinguishing between groups exhibit a greater mean decrease of the Gini coefficient compared to others.

### Integration of CSF and Plasma data sets

To integrate the information from both the plasma and CSF datasets, combined variables were computed. First, Pearson’s correlation analysis was performed between the Ct values of the eleven miRNAs reliably detectable in CSF (Ct_CSF_) and the Ct values of the corresponding miRNAs detected in plasma (Ct_plasma_). A two-tailed p-value cut-off was applied, and alpha was set to 0.05. All significant combinations were used to form new variables by multiplying the corresponding Ct values (Ct_multiplied_). This generated 20 new variables, which were subsequently used to create a heatmap, perform t-tests and PCA and generate LASSO and RF models.

### Identification of discriminatory miRNAs

In a final step, we wanted to select those miRNAs that most accurately discriminated between groups. This feature selection was performed for each of the analyses using the following criteria: 1) p-values resulting from the unpaired t-test were sorted in ascending order and the top five miRNAs with the smallest p-value were selected; 2) the loading scores of PC1 were assessed and the top five miRNAs were selected; 3) the five most influential miRNAs from the RF model were selected based on the mean decrease of the Gini coefficient; 4) the LASSO regression model automatically selects discriminatory features through shrinkage and elimination of less relevant variables by introducing penalties; in this case, three miRNAs were selected. As the CSF data set alone had proven to not efficiently discriminate between groups, these selection steps were only applied to the Ct_plasma_ and Ct_multiplied_ data sets.

### Target prediction enrichment analysis

We used the miRDB database [23] to predict gene targets of selected miRNAs. Next, for each miRNA we selected the top 100 genes sorted by Target score and performed Gene Ontology (GO) enrichment analysis in R using the ontology class “*biological processes*”, the database org.HS.eg.db and the clusterProfiler package [24].

## Results

### Detection of miRNA Expression Patterns in Plasma and CSF

In plasma, a total of 58 miRNAs passed our selection criteria. 23 miRNAs showed stable expression in 29/30 individuals while the remaining 35 miRNAs were detected in all 30 individuals. When visualizing log2fc values using a heatmap, the group of LRRK2_MC_, which included both symptomatic and asymptomatic mutation carriers, showed similar expression patterns and could be distinguished from the sPD group (Fig. 2A). When using the calibrated Ct_plasma_ data, the expression patterns became less apparent but still noticeable (Fig. S5A). In contrast, in CSF only eleven miRNAs survived our selection process, with five miRNAs detected in all 31 individuals and six miRNAs in 30/31 individuals. Expression levels in CSF were generally lower compared to plasma. In CSF, no clear clustering was observable (Fig. 2B). Heatmaps based on calibrated Ct_CSF_ also did not show any clear observable group-specific clusters (Fig. S5B).

**Figure 2:**
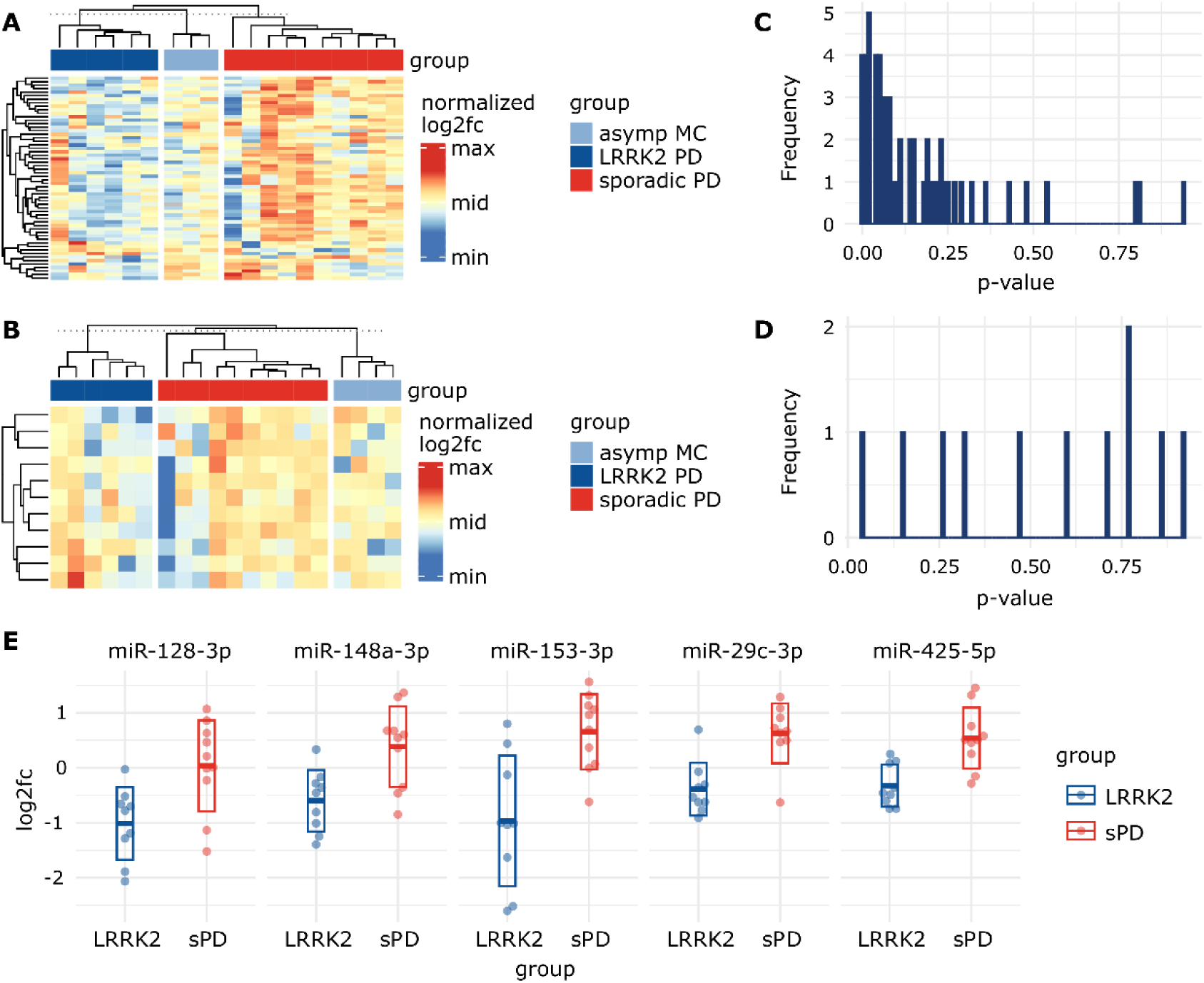
Visualizing miRNA signatures and group differences. **A:** Heatmap based on log2fc values from the plasma dataset, normalized per row. Rows represent different miRNAs while columns represent patients. Grey cells represent missing values. Column sorting function was inactivated to not interfere with group arrangement. The similarity indicated by the column dendrograms is not proportional between groups. **B:** Heatmap based on log2fc values from CSF dataset, normalized per row. This heatmap includes a smaller amount of miRNAs (represented in rows) as they were excluded from the CSF dataset due to missing values. **C:** Histogram showing the distribution of uncorrected p-values obtained from the plasma and the **D:** CSF datasets after performing unpaired t-tests using log2fc values. **E:** Scatterplots display log2fc values from sPD patients and LRRK2_MC_. Five miRNAs with the lowest p-value in the t-tests were selected. Thick line indicates mean while the box indicates the standard deviation.

### Identification of single discriminatory miRNAs through t-test

After performing unpaired t-test on log2fc values in plasma, a total of 17 miRNAs were below a p-value threshold of 0.05 (Table S4). After correction for multiple comparisons by adjusting the p-value threshold to 0.0009, miR-29c-3p showed a significant difference (LRRK2_MC_: -0.39 (SD: ±0.48); sPD: 0.62 (SD: ±0.54), t(56) = 4.21, p = 0.0007). ROC analysis of miR-29c-3p revealed a sensitivity of 90% and specificity of 90% with an AUC of 0.86 (Fig. S6). In CSF, only miR-223-3p showed a p-value below 0.05 (LRRK2_MC_: -0.49 (SD: ±1.05); sPD: 0.62 (SD: ±1.20), t(9) = -2.2, p = 0.041) but due to correction for multiple testing did not pass the p-value threshold of 0.0045.

P-values in the plasma data set were not equally distributed but concentrated at the lower end of the scale (Fig. 2C), while the p-values in the CSF data set appeared to be randomly distributed (Fig. 2D). This indicates that the miRNA expression levels measured in CSF alone seem to contain little information on group discrimination, while in plasma a subset of miRNAs could be used to identify LRRK2_MC_. As a consequence, in the following analyses we primarily focused on the plasma dataset. We sorted p-values from the plasma data set in ascending order and identified the top 5 miRNAs with the lowest p-value (Fig. 2E). These included miR-128-3p (log2fc in LRRK2_MC_: -1.01 (SD: ±0.66); sPD: 0.03 (SD: ±0.83), t(56) = 3.06, p = 0.0071), miR-148a-3p (LRRK2_MC_: -0.60 (SD: ±0.56); sPD: 0.38 (SD: ±0.73), t(56) = 3.31, p = 0.0042), miR-153-3p (LRRK2_MC_: -0.97 (SD: ±1.19); sPD: 0.65 (SD: ±0.69), t(56) = 3.58, p = 0.0035), miR-29c-3p (LRRK2_MC_: -0.39 (SD: ± SD: ±0.48); sPD: 0.62 (SD: ±0.54), t(56) = 4.21, p = 0.0007) and miR-425-5p (LRRK2_MC_: -0.33 (SD: ±0.38); sPD: 0.54 (SD: ±0.55), t(56) = 4.01, p = 0.0010).

### Detecting patient clusters using plasma miRNA expression levels and PCA

PCA explained 82% of the overall variance in the plasma dataset. PC1 (77.4% of variance) explained the majority of the variance and appeared to discriminate groups, while PC2 only explained 4.6% of the overall variance (Fig. 3A). Interestingly, all LRRK2_MC_ clustered together while the various *LRRK2* mutations could not be distinguished (Fig. 3A). PC1 discriminated LRRK2MC from sPD with a sensitivity of 100% (9/9) and a specificity of 70% (7/10). PC1 and PC2 combined reached the same sensitivity with a specificity of 80% (8/10) (Fig. S7A). Next, we performed PCAs including the healthy controls (Fig. 4). While clustering was less clear when comparing HC to sPD, for comparison of HC and LRRK2_MC_ clear clustering was observable (Fig. 4A and 4B). The clearest clustering was achieved when comparing LRRK2_MC_ with sPD (Fig. 4C). When including all three groups in the PCA, clustering becomes less clear, yet LRRK2_MC_ remains notably separated from sPD and HCs (Fig. 4D). The PCA of the CSF data explained 84.2% of the overall variance, with both PC1 (75.4% of variance) and PC2 (8.8% of variance) not able to differentiate between the groups (Fig. 3B).

**Figure 3:**
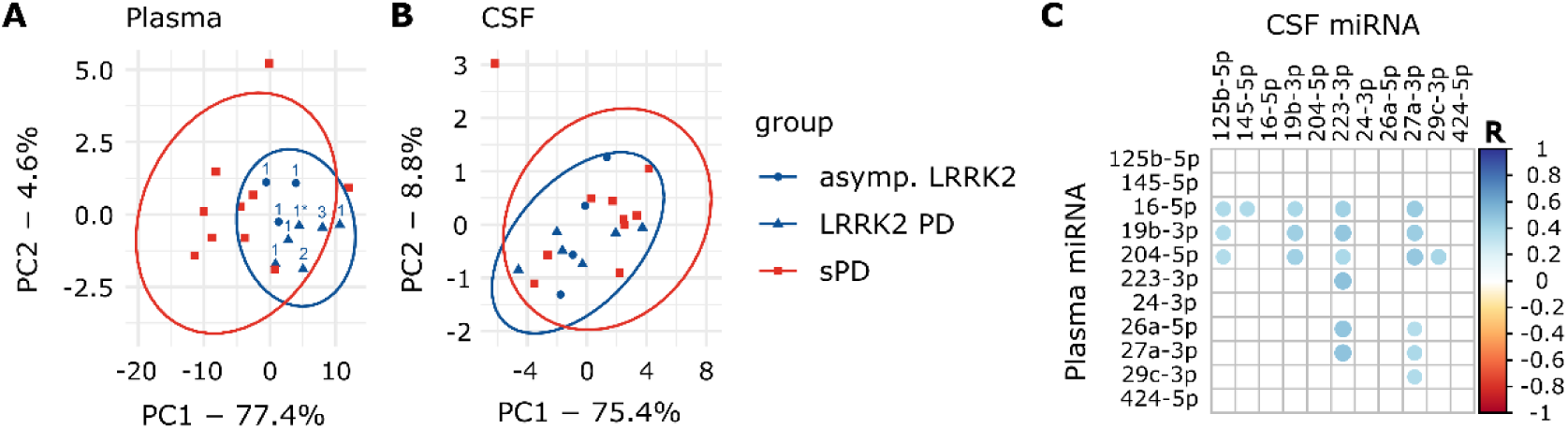
PCA graphs and correlation of plasma and CSF data. **A:** PCA graph from plasma and **B:** CSF Ct values. Ellipses indicate 95% confidence interval. Numbers indicate *LRRK2* mutation (1: G2019S, 1*: G2019S + G1819, 2: R1441C, 3: I2020T). In the plasma PCA graph, group ellipses overlap while the blue LRRK2_MC_ ellipse trends to only include individuals of the LRRK2_MC_ group. Based on PCA, the LRRK2_MC_ group could be interpreted as a subpopulation of all PD patients. **C:** Correlation matrix presenting statistically significant (p < 0.05) correlations between Ct_plasma_ and Ct_CSF_ values of the eleven miRNAs included in the CSF dataset.

**Figure 4:**
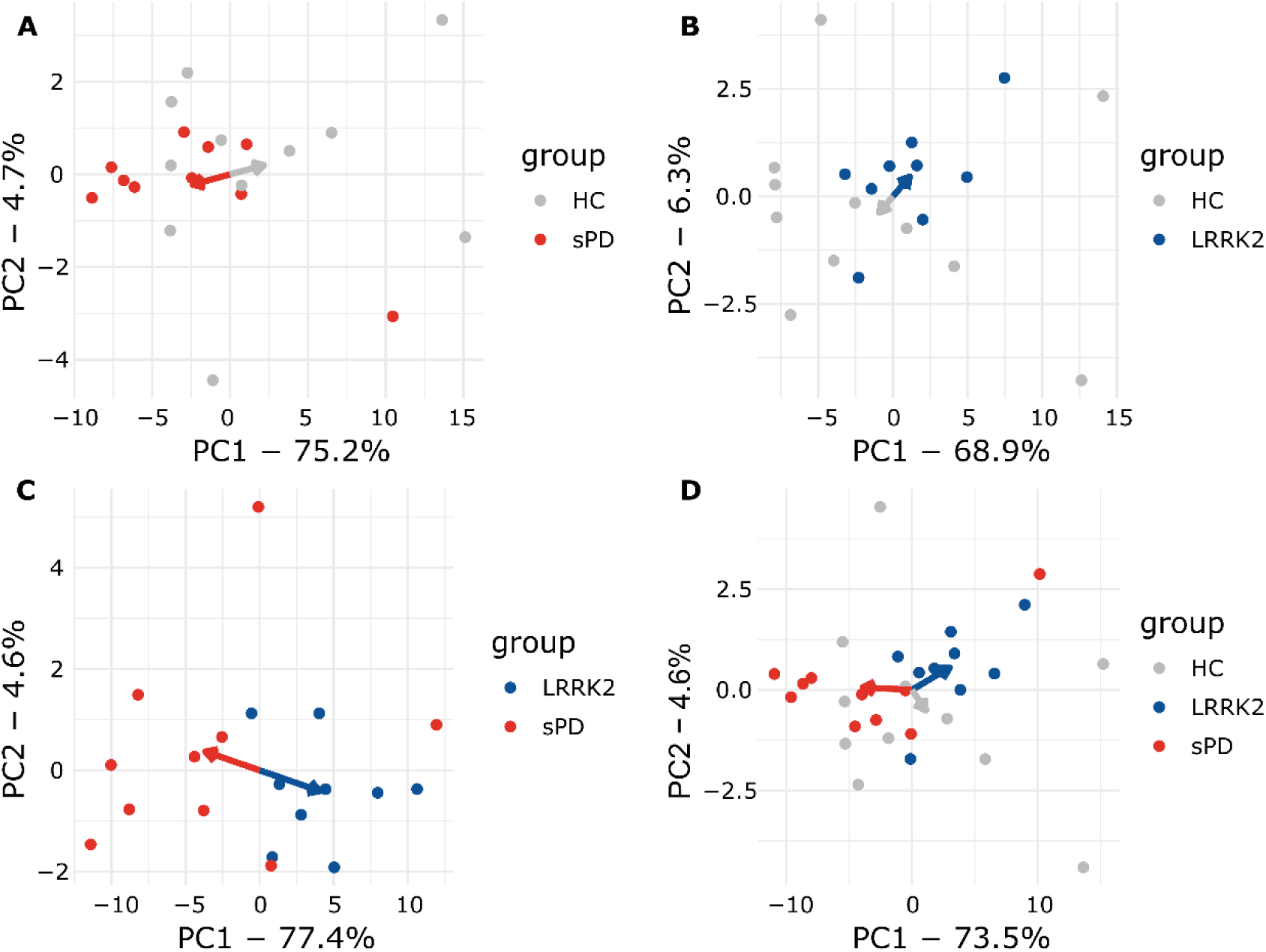
Plasma PCA plots comparing all groups. **A:** sPD and HC**. B:** LRRK2_MC_ and HC. **C:** LRRK2_MC_ and sPD. **D:** All groups. Arrows point towards the respective mean of PC1 and PC2. In **A**, **B** and **C** clear separation can be observed as indicated my arrows pointing towards opposing directions. When including all groups in **D**, clustering is less apparent but still observable.

### miRNA expression levels in CSF correlate to Plasma levels

A total of 20 miRNA combinations from Ct_plasma_ and Ct_CSF_ values showed significant correlation (Fig. 3C) with R-values ranging from 0.36 to 0.51 (Table S5). Significantly correlating combinations were used to obtain new variables by multiplication (Ct_multiplied_).

### Multiplied Ct values discriminate groups in t-test and PCA

When creating a heatmap based on the Ct_multiplied_ data, no clear group-specific clusters were noticeable (Fig. S8A). In the t-test ten miRNA-combinations reached a p-value below 0.05 (Table S6). The combination of miR-29c-3p from plasma and miR-27a-3p from CSF passed the adjusted p-value threshold of 0.0025 (LRRK2_MC_: 930.8 (SD: ±61.6); sPD: 813.6 (SD: ±55.9), t(18) = 4.2, p = 0.0007). PCA for Ct_multiplied_ discriminated the groups while explaining 90.2% of the total variance (PC1: 83.2%, PC2: 7%) (Fig. S8B). Groups were separated based on PC1 with a sensitivity of 100% (9/9) and a specificity of 80% (8/10). Combining PC1 and PC2 did not improve sensitivity or specificity (Figure S7B).

### LASSO Regression and Random Forest Models predict group membership

LASSO regression and the RF model were performed using the Ct_plasma_ and the Ct_multiplied_ dataset (Fig. 5A). Sensitivity of the LASSO model was 88.8% (8/9) for both Ct_plasma_ and Ct_multiplied_. Specificity was 70.0% (7/10) (Ct_plasma_) and 80.0% (8/10) (Ct_multiplied_), respectively (Fig. 5B). Predictions from the RF models using Ct_plasma_ had a mean sensitivity of 84.2% (95% CI: 83.1% – 85.3%) and a mean specificity of 70.1% (95% CI: 69.9 – 70.2%). When using Ct_multiplied_ mean sensitivity was 73.6% (95% CI: 72.5% – 74.6%) and mean specificity was 80 % (95% CI: 80% – 80%) (Figure 5B). When comparing the two models using an unpaired t-test, sensitivity was significantly better when using Ct_plasma_ (t(98) = -13.8, p<0.001), while specificity was higher when using Ct_multiplied_ (t(98) = 99, p<0.001). Classifications performed by the RF model were further analyzed by looking into proximity scores of patient pairs. The resulting heatmap for the Ct_plasma_ dataset indicated that patients within the same group exhibit greater proximity to each other than patients from different groups (Fig. 5C). This becomes even more apparent in the model using Ct_multiplied_ (Fig. 5D). Next, mean decrease of Gini scores were calculated for both of these RF models (Fig. 5E and 5F) to assess variable influence on classification accuracy. Interestingly, in both RF models, miR-223-3p had the highest score, indicating a great impact on the model performance. This miRNA had already been observed in the plasma t-test, where it indicated a potential for group discrimination with an uncorrected p-value of p=0.051.

**Figure 5:**
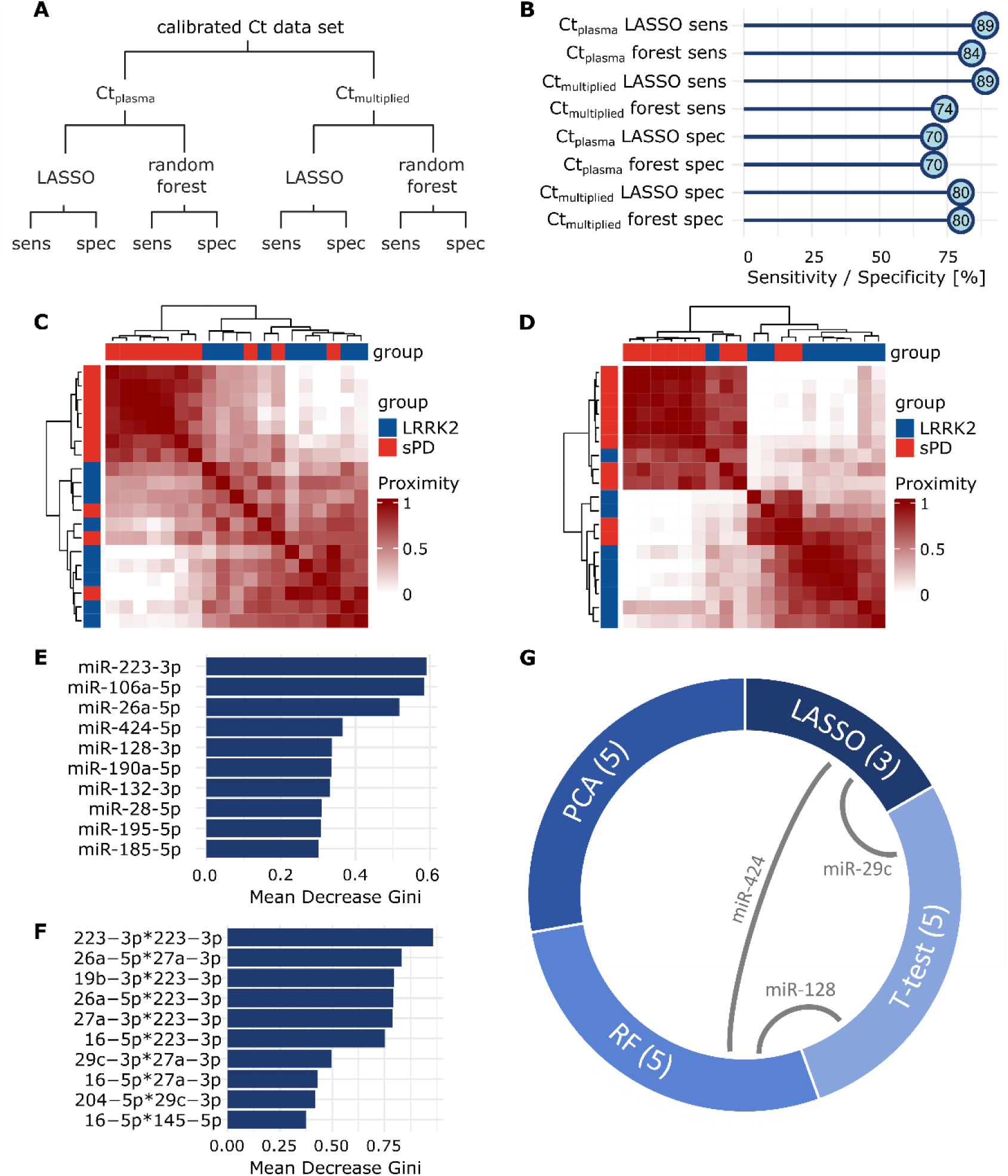
Group prediction with LASSO and Random Forest. **A:** Overview of performed analyses and respective read-outs. **B:** Sensitivity and specificity values acquired by LASSO regression and Random Forest, using Ct_plasma_ and Ct_multiplied._ **C:** Heatmaps display the proximity scores calculated in RF for all sPD patients (red) and LRRK2_MC_ (blue) using Ct_plasma_ or **D:** Ct_multiplied_. Groups clearly separate along mutation status. **E:** Mean Decrease Gini scores were calculated after building the respective RF models for both the Ct_plasma_ and **F:** the Ct_multiplied_ data set. Values were sorted in descending order and the top ten miRNAs are displayed. High decrease of the Gini coefficient translates to a high impact on the performance of the respective RF model. **G:** From each analysis, most influential or discriminatory miRNAs were extracted. Chord diagram displays the overlap of miRNAs selected from the different analyses. Numbers in brackets display set size and grey connecting lines indicate miRNA overlaps.

### Selection of discriminatory miRNAs identified by multiple tests or models

Finally, for each test or model, we identified those miRNAs, that showed the greatest potential for group separation or prediction (Table 2). From the RF models, PCA and the t-tests, we extracted five miRNAs each, while the LASSO model identified three miRNAs. Interestingly, of the selected miRNAs, three were identified by more than one method; 1) miR-29c-3p was identified in t-tests and LASSO, 2) miR-128-3p in t-test and RF, and 3) miR-424-5p in LASSO and RF (Fig. 5G, Fig. S9). GO analysis was performed on the four miRNAs and we identified associated biological functions and processes (number of significant annotations identified: miR-223-3p: 25, miR-29c-3p: 60, miR-128-3p: 0, miR-424-5p: 3, see Figure S10). Interestingly, the annotations for miR-223-3p included neuron death, regulation of neuron death and neuron apoptotic process.

**Table 2:** miRNAs selected as most influential by the different models.

| t-test selected | PCA selected | LASSO selected | Random forest selected |
| --- | --- | --- | --- |
| **miR-29c-3p** | miR-19b-3p | **miR-29c-3p** | miR-26a-5p |
| **miR-128-3p** | miR-27a-3p | **miR-424-5p** | miR-106a-5p |
| miR-148a-3p | miR-28-5p | miR-532-5p | **miR-128-3p** |
| miR-153-3p | miR-30b-5p |  | *miR-223-3p* |
| miR-425-5p | miR-185-5p |  | **miR-424-5p** |

The table displays miRNAs that were identified as relevant from each model or test. miRNAs highlighted in bold were identified in more than one analysis. Of note, miR-223-3p (italic) was extracted from both the Ct<sub>plasma</sub> and the Ct<sub>multiplied</sub> RF models.

## Discussion

In this proof-of-concept study, we examined the extracellular miRNA signatures in plasma and CSF derived from LRRK2_MC_ and sPD patients for LRRK2-dependent patterns. We discovered that plasma miRNA expression levels can distinguish sPD from LRRK2_MC_. MiRNAs have been extensively studied in PD, but to the best of our knowledge, this is the first study to use machine learning and a broad data set of miRNA expression levels to distinguish genetic from sporadic PD.

Our results show that PCA separated LRRK2_MC_ from most individuals with sPD without differentiating between the various *LRRK2* mutations. Separation was also observable in PCA comparing LRRK2_MC_ and HC as well as in an analysis comparing all three groups. This indicates that the LRRK2 mutation status has a measurable effect on the extracellular miRNA signature in LRRK2_MC_. Some sPD individuals clustered in proximity to the LRRK2_MC_ individuals, raising the interesting possibility that this method could be used to identify patients who would benefit from a LRRK2 targeted therapy. However, this question needs to be addressed using larger cohorts in the context of LRRK2-inhibitor trials. Regular assessment of the identified miRNA signatures could help monitor treatment efficacy in therapies targeting LRRK2, where a shift in miRNA expression levels could indicate target engagement.

We extracted discriminatory miRNAs from each analysis and found that some miRNAs were overlapping between models, namely miR-29c-3p, miR-128-3p, miR-424-5p and miR-223-3p.

miR-29c-3p has been reported to play a role in disease modulation by mediating neuroinflammation and has been found to be involved in apoptotic processes [25]. It has further been proposed as a PD specific marker in several studies, but contradicting findings on expression levels compared to controls exist: studies report either downregulation [26,27] or upregulation [28] in serum of sPD patients compared to HCs. One study found upregulation in serum of sPD patients, but no dysregulation in LRRK2 associated PD [29].

miR-223-3p has been reported to be upregulated in midbrain dopamine neurons [30] and serum of PD patients [31]. It was shown to be involved in the modulation of inflammasome activity [32]. LRRK2 has long been suspected to play a role in inflammation and e.g., was shown to affect microglial activation and pro-inflammatory cytokine production [33]. Further, the GO annotations identified for miR-223-3p included processes specific to neurodegeneration underlining a possible influence in neurodegenerative disorders such as PD.

miR-128-3p has previously been identified as a potential treatment target due to its ability to protect neurons from apoptosis. The upregulation in the sPD group could therefore be reflective of a compensatory mechanism, that might not be relevant in individuals carrying a *LRRK2* mutation.

miR-424-5p was shown to be increased in the forebrain of PD patients, while being associated with FOXO1 [34]. FOXO1 activity is induced by LRRK2 [35] and thereby links altered miR-424-5p levels to LRRK2 activity.

The low RNA abundance in CSF compared to plasma made miRNA detection difficult, resulting in the exclusion of many miRNAs from CSF analyses. This reduced the complexity of miRNA signatures we could assess in CSF, which may explain the poor performance of CSF-based analyses. An alternative explanation could be that LRRK2 protein expression levels are known to be low in the CNS [4,5] despite their apparent relevance in the pathophysiology of PD. In contrast, peripheral organs such as the lung and the kidney display higher expression levels, which could also explain the increased accuracy of models based on Ct_plasma_ miRNA profiles in identifying LRRK2_MC_. We have further analyzed the relation of miRNA expression levels in CSF and plasma and found that for most miRNAs, these two biofluids seem to display very different signatures. However, through correlation analysis we have identified a subset of miRNAs whose expression levels in the CNS and the periphery seem related. We convoluted correlated miRNA data sets in order to test whether Ct_multiplied_ would provide RF models with significantly higher sensitivity or specificity, which was not the case. While RF models based on Ct_multiplied_ still performed well and slightly different than RF models based on Ct_plasma_, when facing the discussed classification problem there seems to be no clear benefit from adding CSF data to the model.

In summary, this proof-of-concept study showed promising results and deepens our understanding of LRRK2-associated PD, but it has limitations we want to address. First, the number of individuals included in the present study is relatively small. When utilizing group sorting algorithms for classification, larger sample sizes improve robustness and replicability. Ideally, the data set should be divided into a training and a testing data set to avoid overfitting. Additionally, a completely independent data set should then be used to replicate the predictions. While we did perform cross-validation using LOOCV or out-of-bag samples, this cannot fully replace validation in a testing or replication cohort. The results therefore have to be considered with caution until replicated in larger cohorts. Further, we selected miRNAs to be included in our qPCR panel based on the literature and therefore only considered miRNAs that have already been reported as dysregulated in PD or other neurodegenerative diseases. This may have introduced a bias and preventing discovery of novel contributing miRNAs. Finally, we found no evident advantage to employing multi-layered or composited readouts over standard t-tests. While we still believe that in diseases as complex as PD basing classifications on multiple variables is an interesting approach, this concept has still to be improved and repeated in larger and more complex cohorts.

## Conclusion

In conclusion, in this proof-of-concept study we showed that LRRK2 mutation status impacts the extracellular miRNA signature measured in plasma and shows promise to separate LRRK2_MC_ from sPD. Monitoring changes of the extracellular miRNA signatures upon e.g. LRRK2 inhibition could be used to study drug efficacy or target engagement. The potential of multi-layered approaches trying to identify sporadic PD patients with a relevant role of LRRK2-related pathways are another potential application which needs to be thoroughly assessed in larger cohorts.

## List of Abbreviations

CSF: Cerebrospinal fluid
Ct_CSF_: Ct values from CSF dataset
Ct_multiplied_: multiplied Ct values from CSF and plasma dataset
Ct_plasma_: Ct values from plasma dataset
Fc: Fold change
GO: Gene Ontology
HC: Healthy control
LASSO: least absolute shrinkage and selection operator
LEDD: levodopa equivalent daily dose
Log2fc: log_2_(Fold change)
LOOCV: Leave-One-Out Cross-Validation
LRRK2: Leucine-Rich Repeat Kinase 2
LRRK2_MC_: *LRRK2* mutation carrier
miRNA: microRNA
MoCA: Montreal Cognitive Assessment
PCA: Principal component analysis
PD: Parkinson’s disease
qPCR: real-time quantitative PCR
RF: Random Forest
RT: Reverse transcription
sPD: sporadic PD
UPDRS: Unified Parkinson Disease Rating Scale

## Declarations

### Ethics approval and consent to participate

The study was approved by the Hertie Institute for Clinical Brain Research Biobank and the ethics committee of the medical faculty of the University of Tübingen and University Clinic Tübingen (project ID: 199/2011B01). All participants gave written, informed consent.

### Availability of data and materials

The generated and analyzed data are available from the corresponding author upon reasonable request.

### Authors’ Contributions

FK conceptualized the study. FK and TG supervised the study. LJB performed the experiments and the statistical analyses. LJB generated all figures. LJB and FK drafted a first version of the manuscript. All authors read, edited and approved the final manuscript.

## Acknowledgements

Samples were obtained from the Neuro-Biobank of the University of Tuebingen, Germany (https://www.hih-tuebingen.de/en/about-us/core-facilities/biobank/). This biobank is supported by the local University, the Hertie Institute and the DZNE.

## Supplementary Material

**Figure S1:**
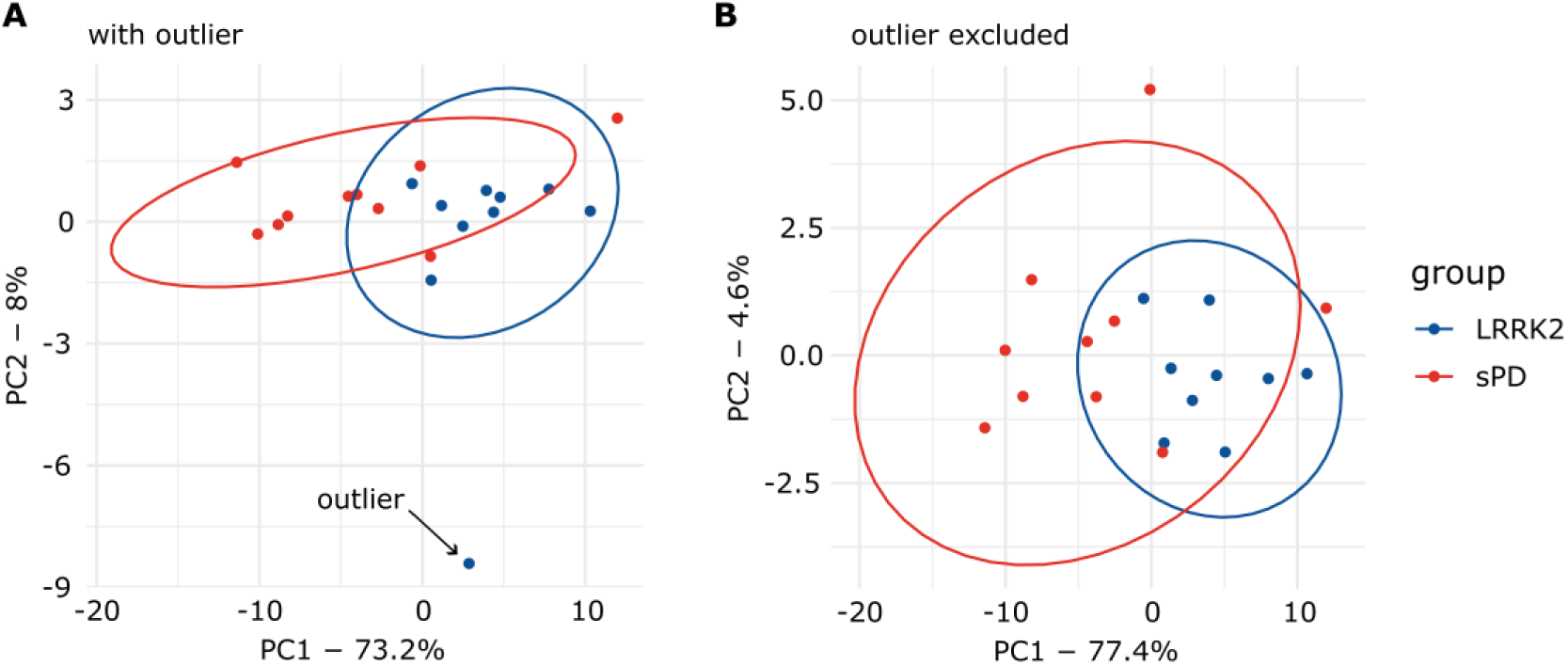
Identification of an outlier via PCA. **A:** The PCA graph on the left displays data points from all sPD patients and LRRK2_MC_. One of the asymptomatic LRRK2_MC_ clearly behaved as an outlier and was therefore excluded from further analysis. **B:** PCA was repeated with the outlier excluded.

**Figure S2:**
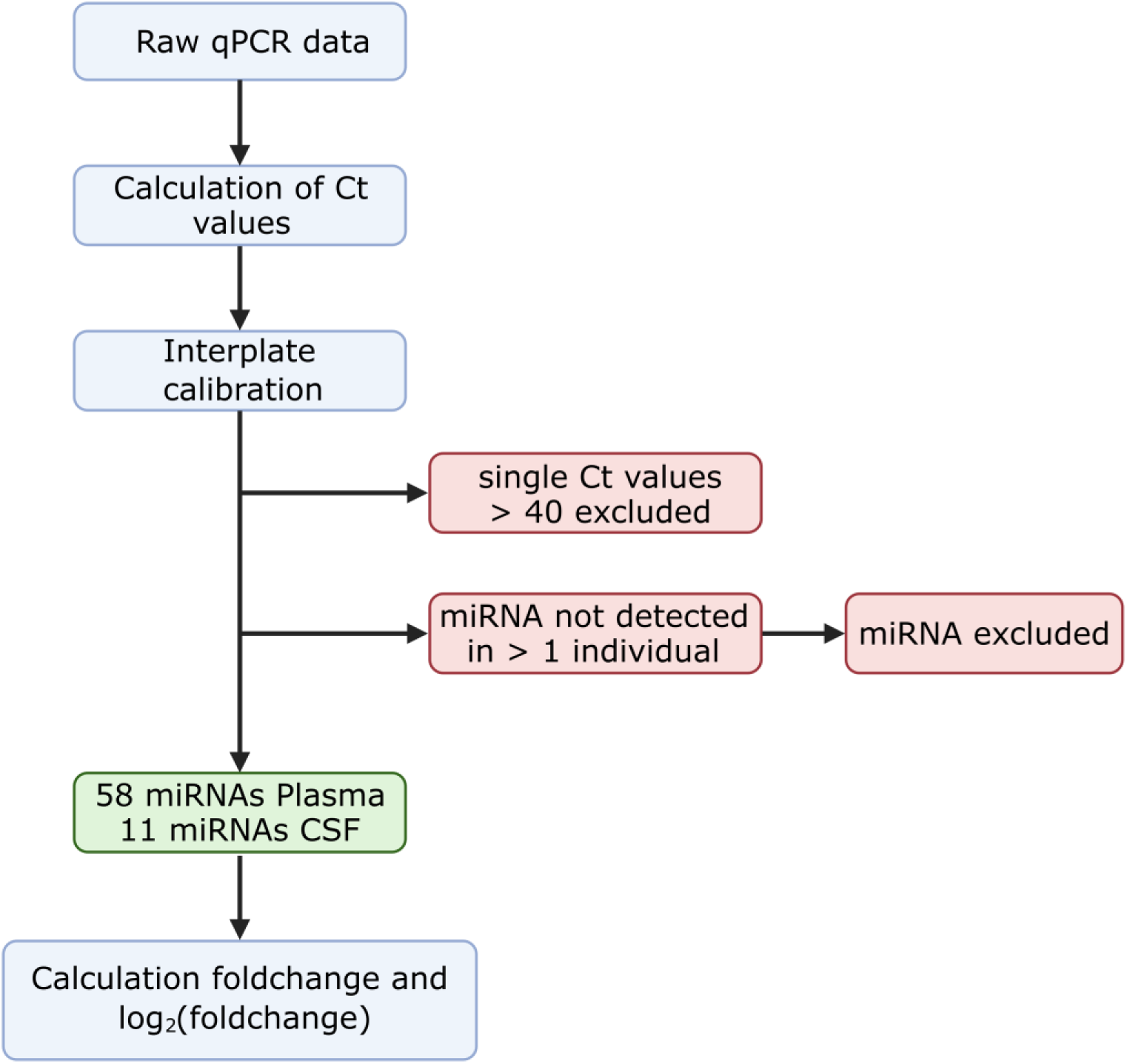
Processing of Raw data. Overview of data processing and selection process. Created with BioRender.com.

**Figure S3:**
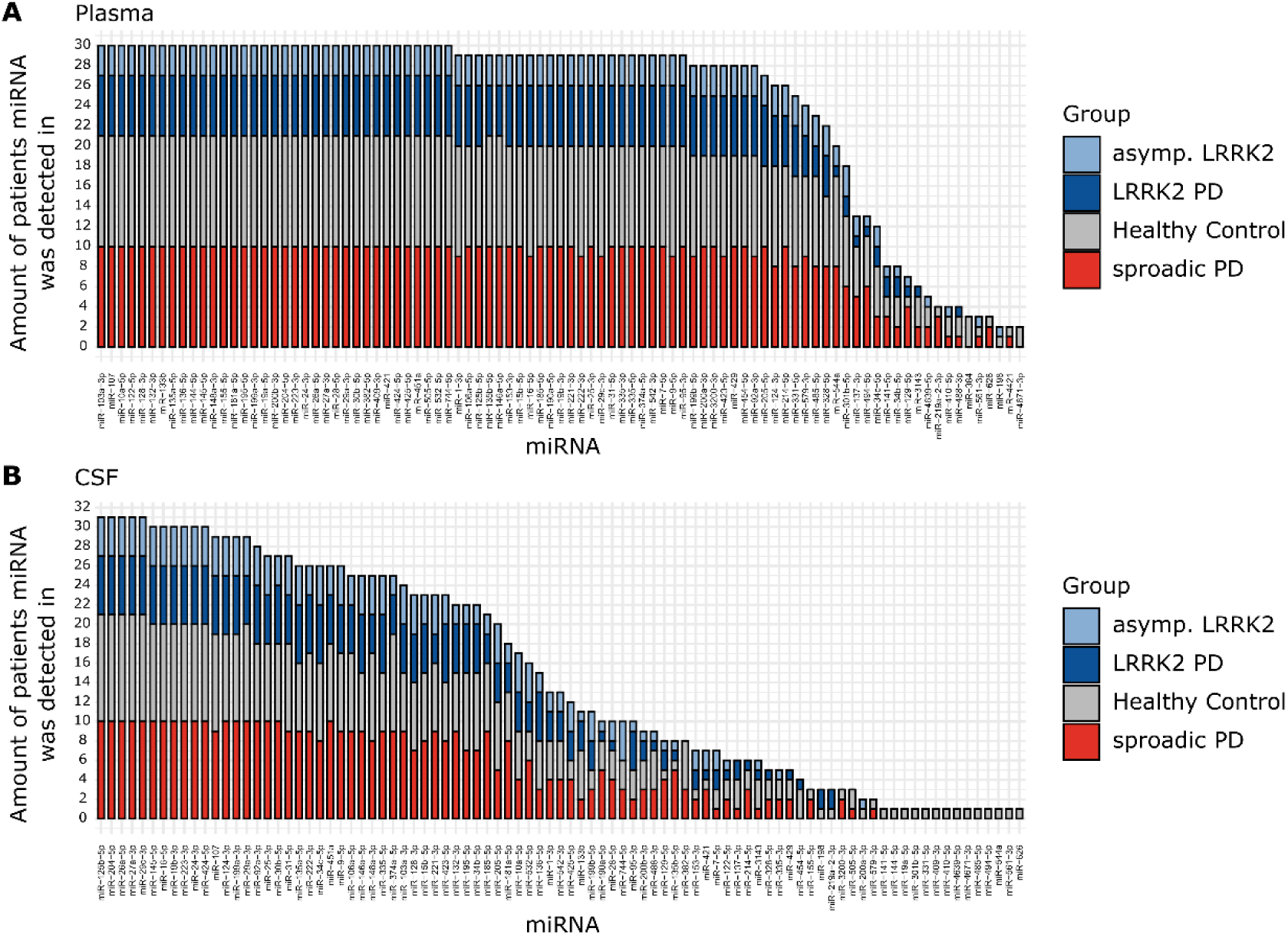
Group-wise Expression Distribution of miRNAs. **A:** The histograms show the number of individuals any given miRNA (x-axis) was found to be expressed in, either in plasma or **B:** CSF. Colors indicate group affiliation.

**Figure S4:**
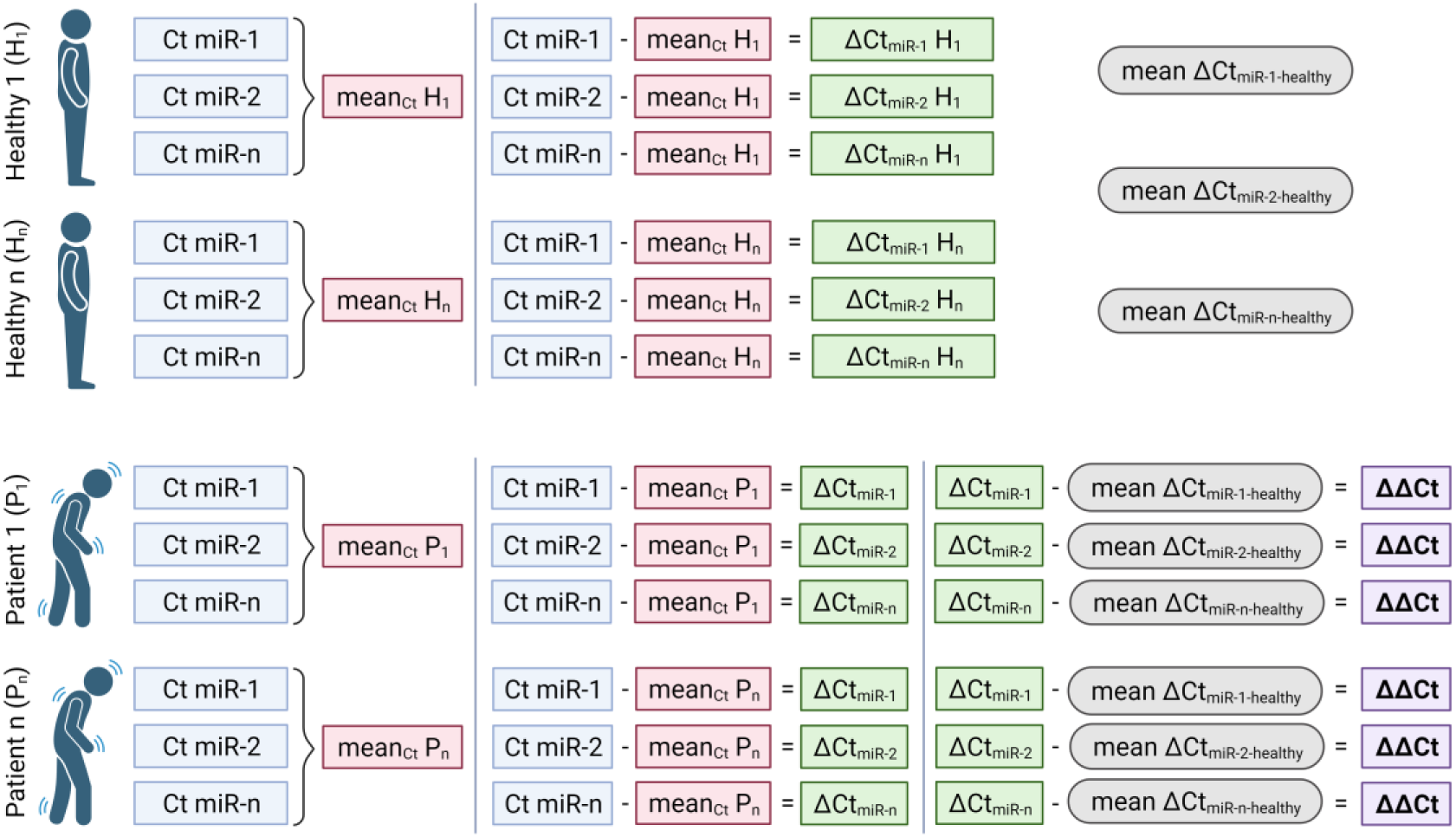
Calculation of ΔΔCt values. Overview of calculation of ΔΔCt values. Created with BioRender.com.

**Figure S5:**
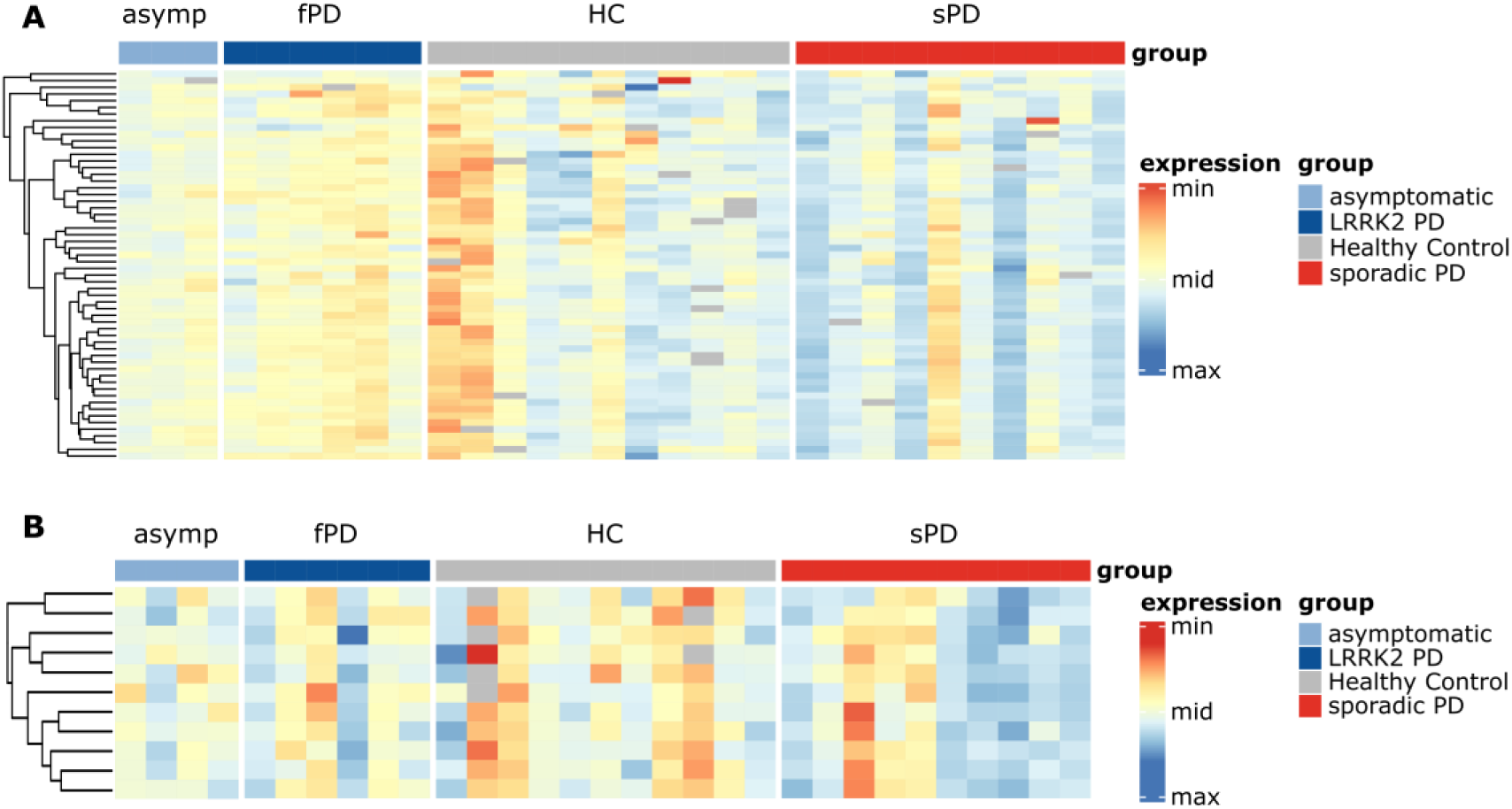
Heatmaps based on Ct values. **A:** Heatmaps for Ct_plasma_ and **B:** Ct_CSF_, normalized per row. Rows represent miRNAs, columns represent individuals. Grey cells represent missing values.

**Figure S6:**
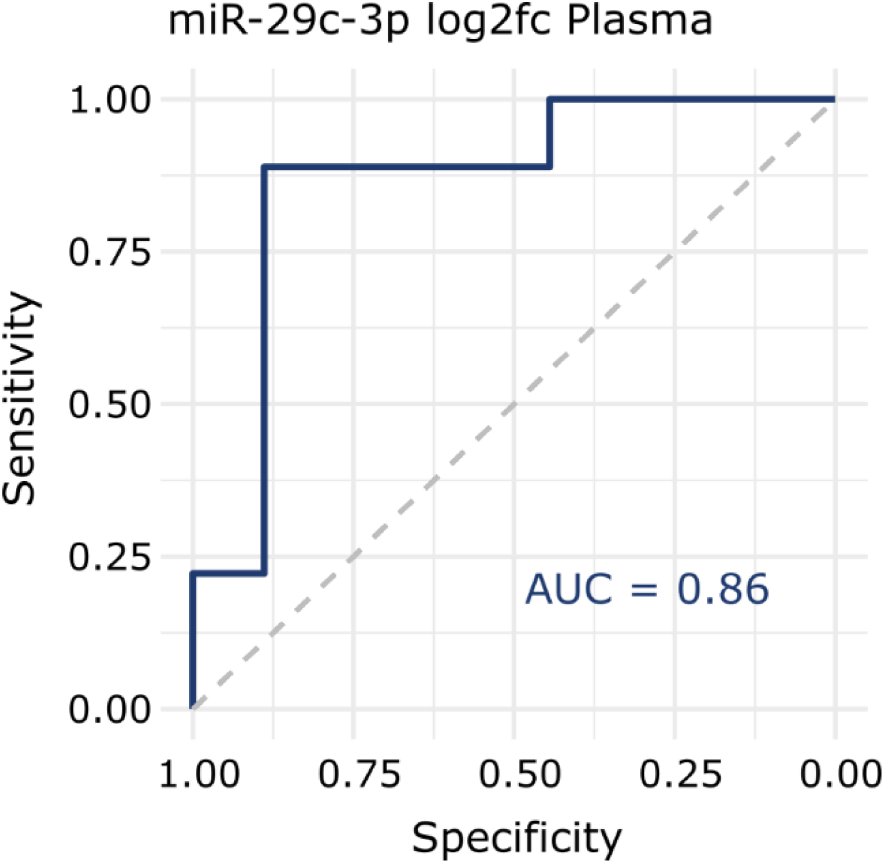
ROC curve for miR-29c-3p. ROC curve displaying the sensitivity and specificity of a classification in LRRK2_MC_ and sPD based on plasma log2fc values of miR-29c-3p.

**Figure S7:**
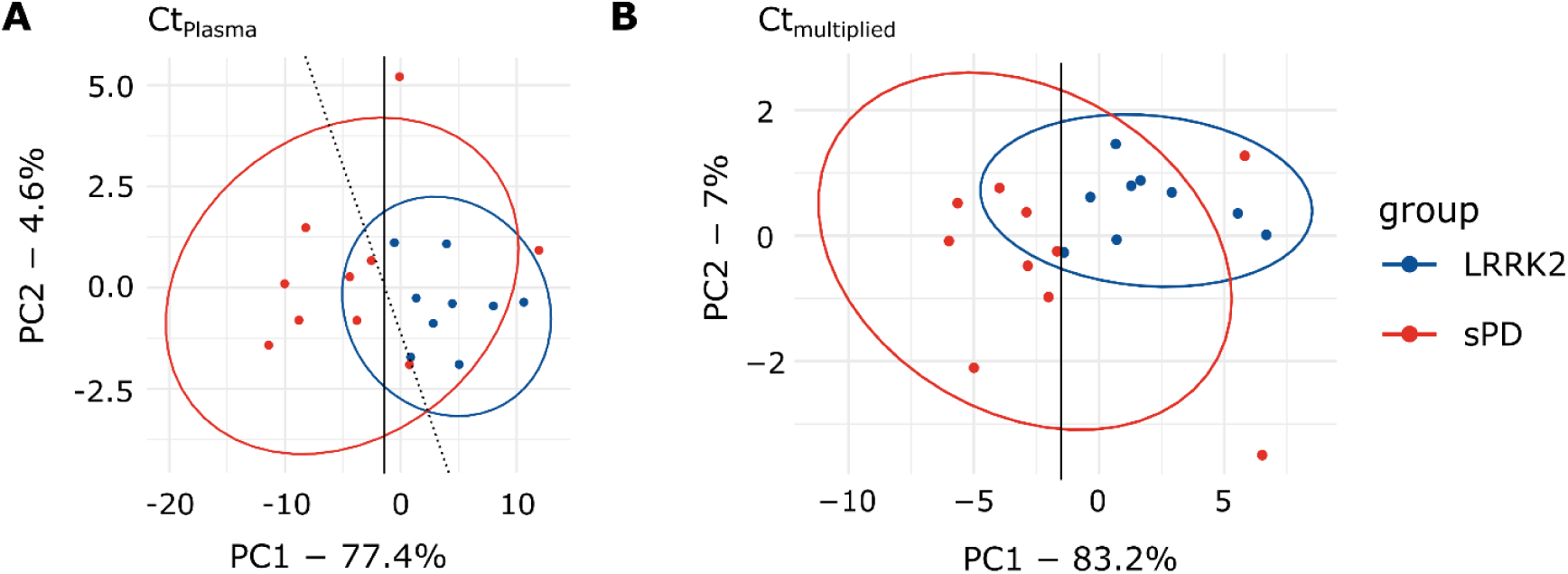
PCA graphs including lines for evaluation of discrimination based on sensitivity and specificity. A: Ct_plasma_. PC1 (solid line) achieves a sensitivity of 100% (9/9) and a specificity of 70% (7/10). PC1 and PC2 combined (dotted line) reach the same sensitivity with a specificity of 80% (8/10). B: Ct_multiplied_. PC1 achieves separation with sensitivity of 100% (9/9) and specificity of 80% (8/10).

**Figure S8:**
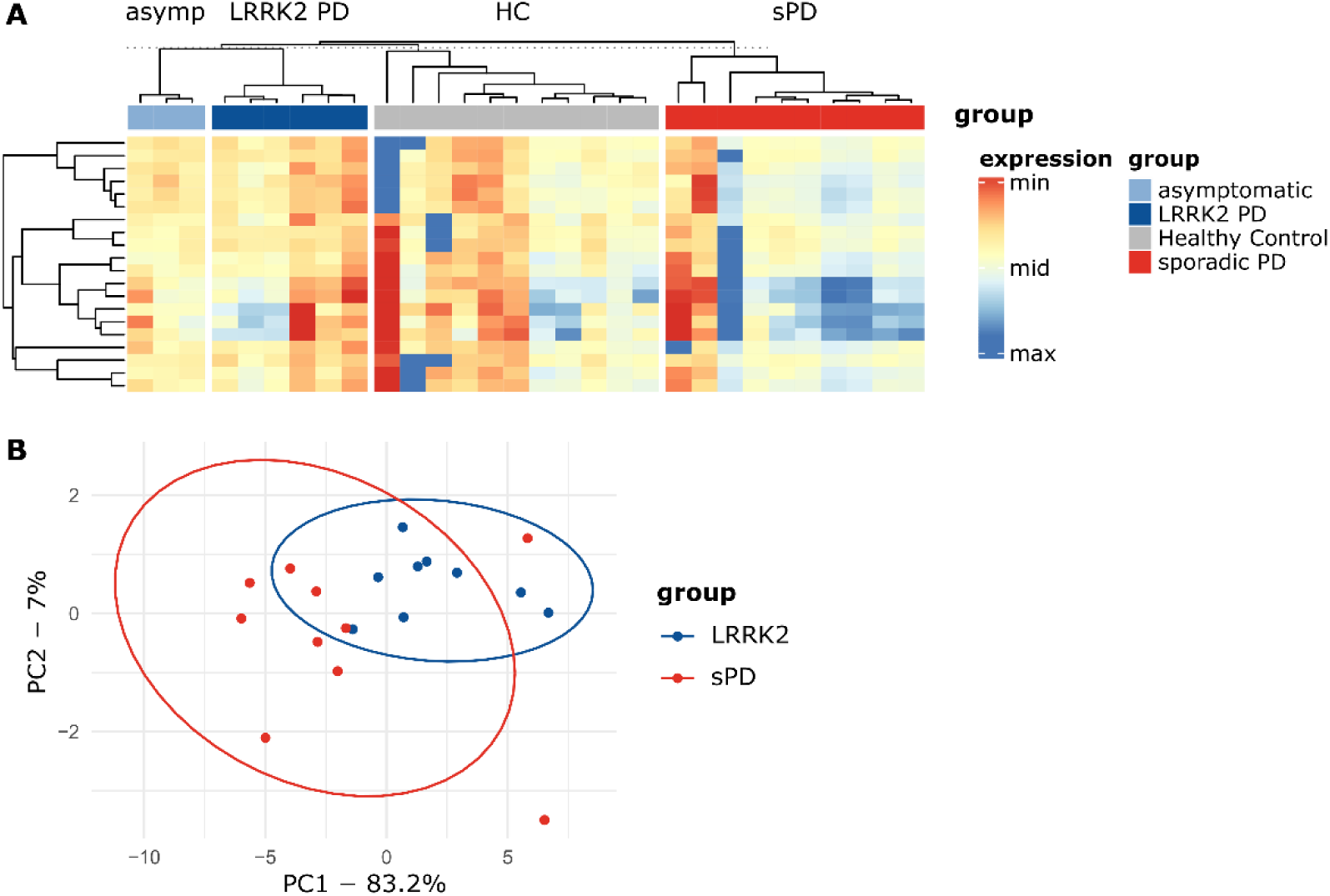
Heatmap and PCA resulting from multiplied variables. **A:** Ct_multiplied_ are plotted as a heatmap for all groups. Again, rows display miRNAs, while columns display individuals. **B:** When performing PCA with Ct_multiplied_, group separation was observable.

**Figure S9:**
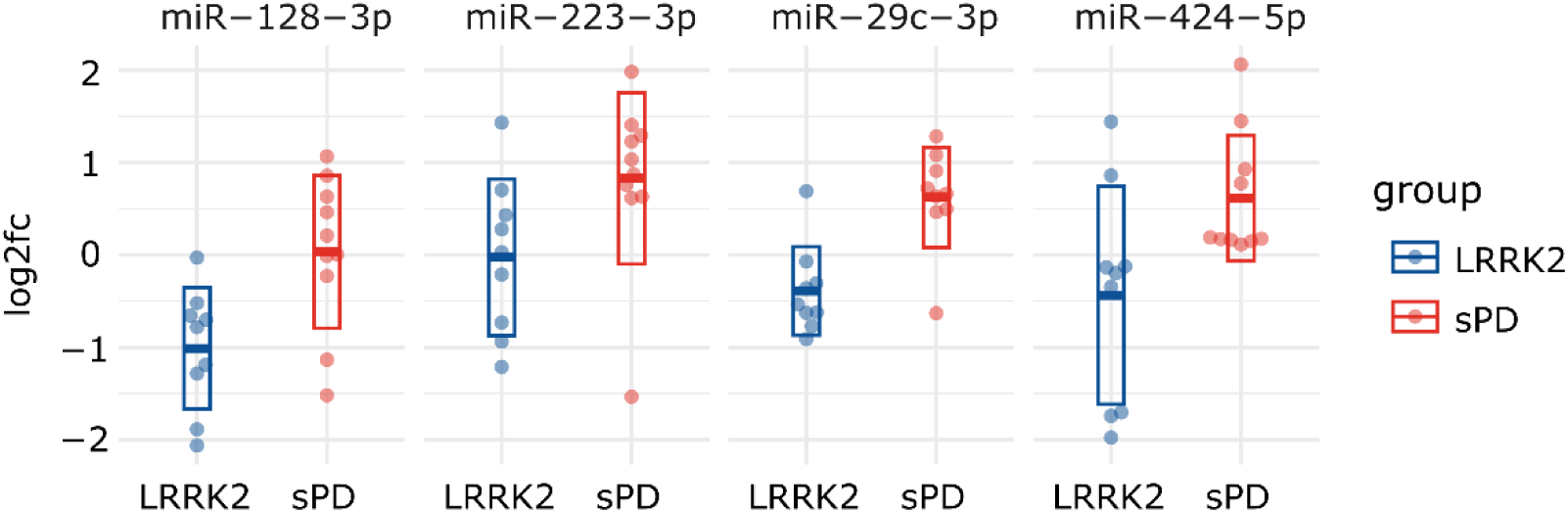
Scatterplots of log2fc values for the four discussed miRNAs. Thick line indicates mean while the box indicates the standard deviation.

**Figure S10:**
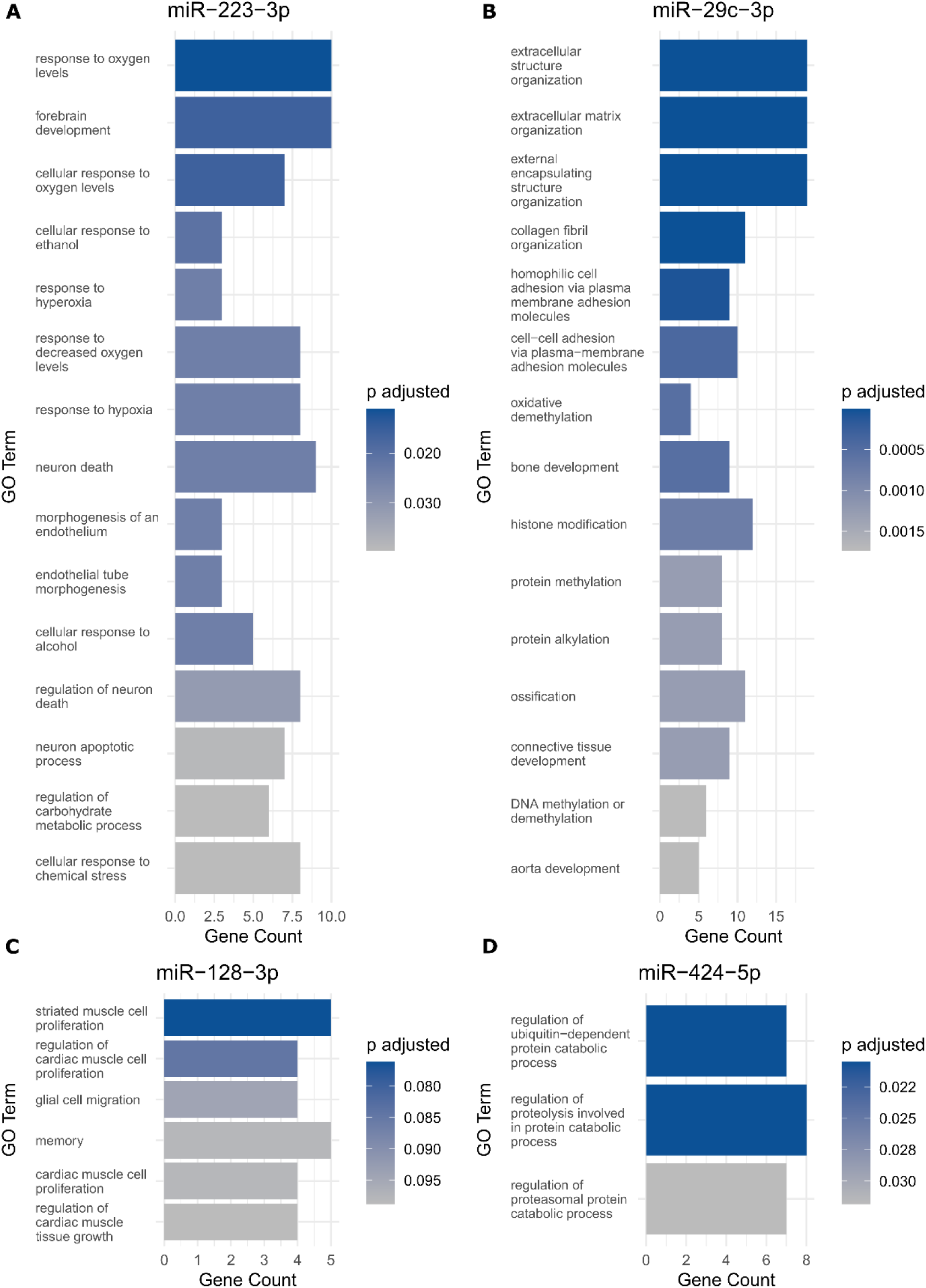
GO analysis of identified miRNAs. **A**: miR-223-3p. 25 significant annotations were identified and sorted by adjusted p-value. Top 15 annotations are displayed. **B**: miR-29c-3p. 60 significant annotations were identified, top 15 annotations are displayed. **C**: miR-128-3p. There were no annotations with an adjusted p-value < 0.05. 6 annotations with an adjusted p-value < 0.01 are displayed. **D**: miR-424-5p. 3 Annotations with an adjusted p-value < 0.05 were identified.

**Table S1:**
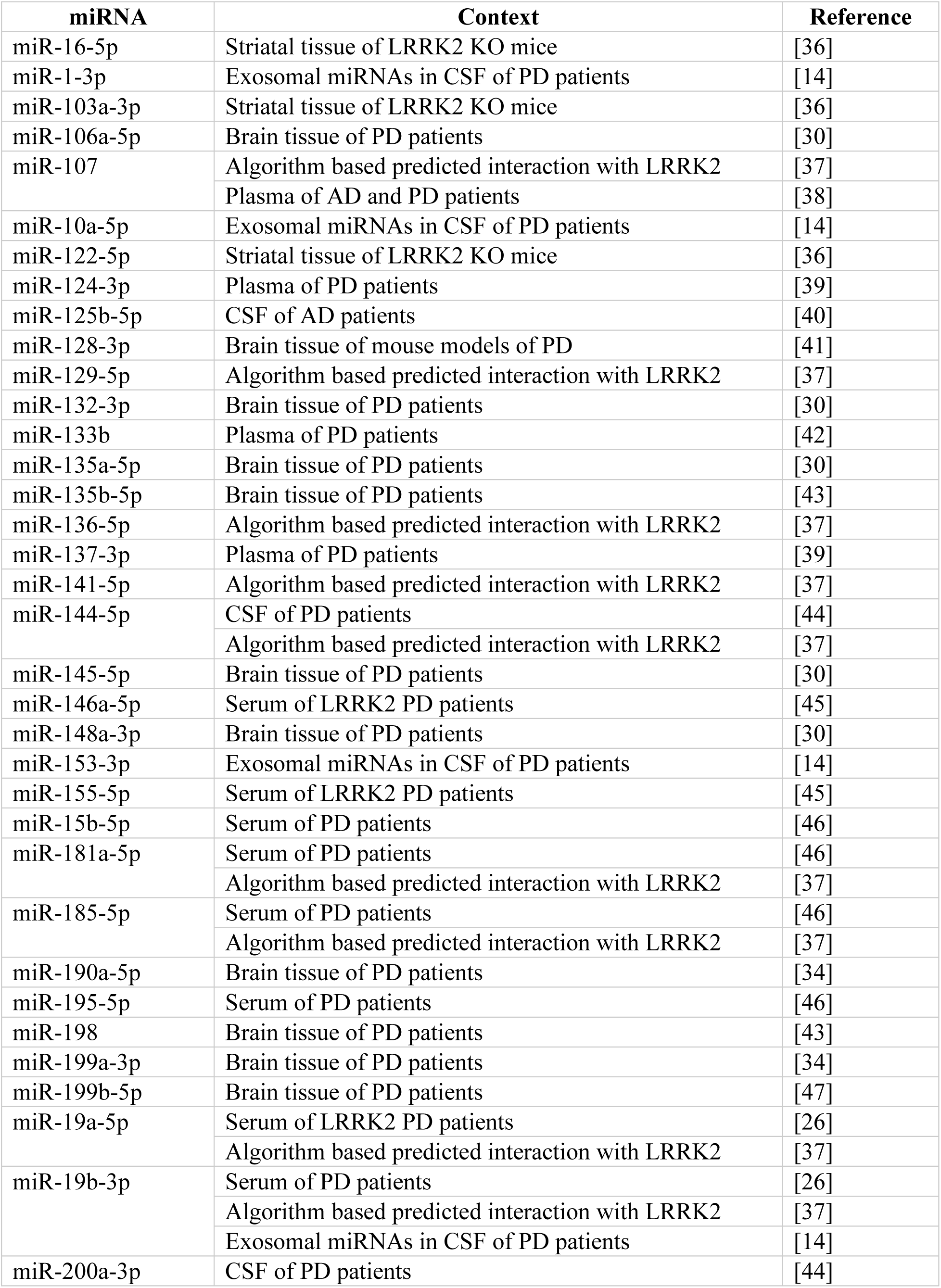
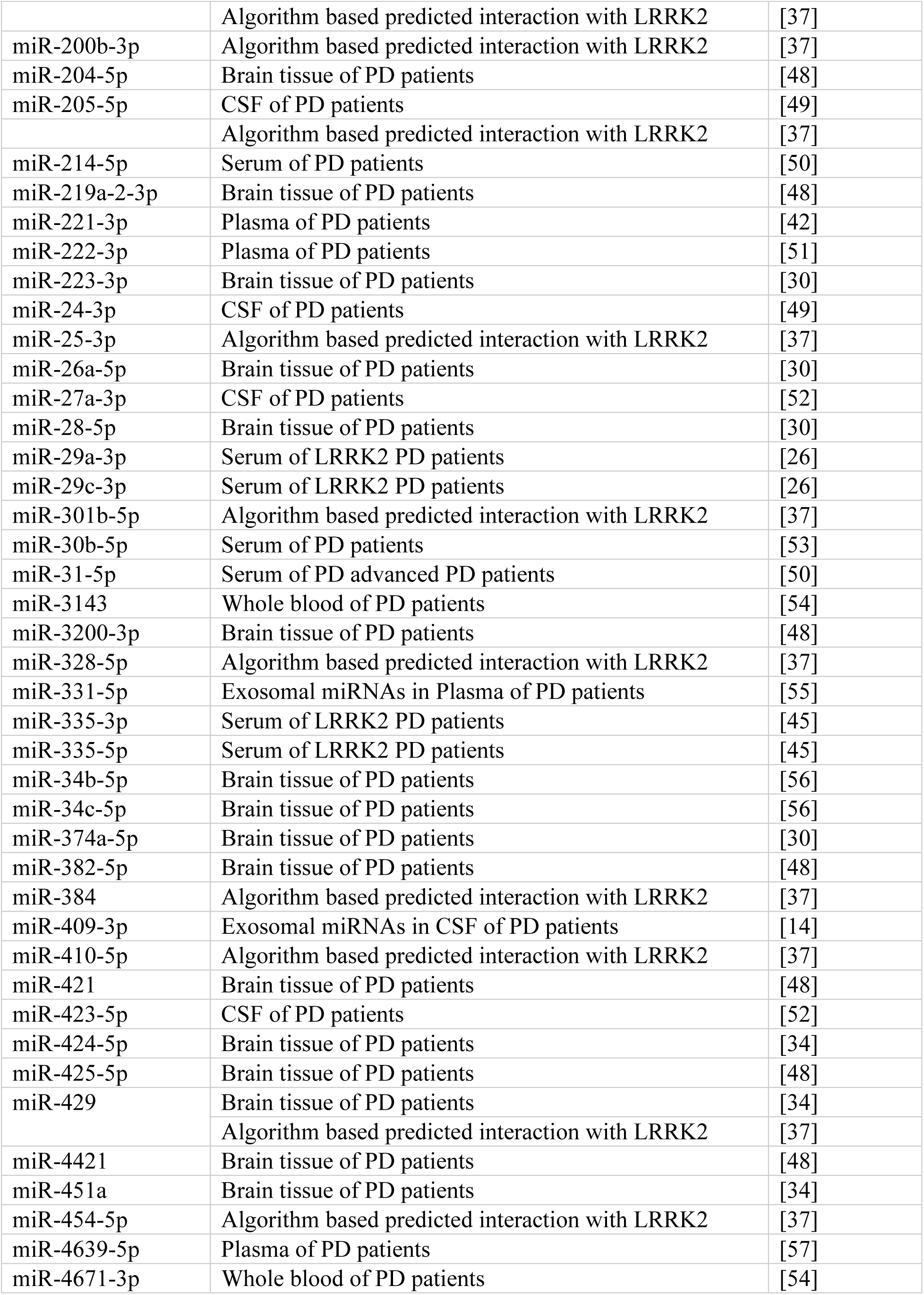
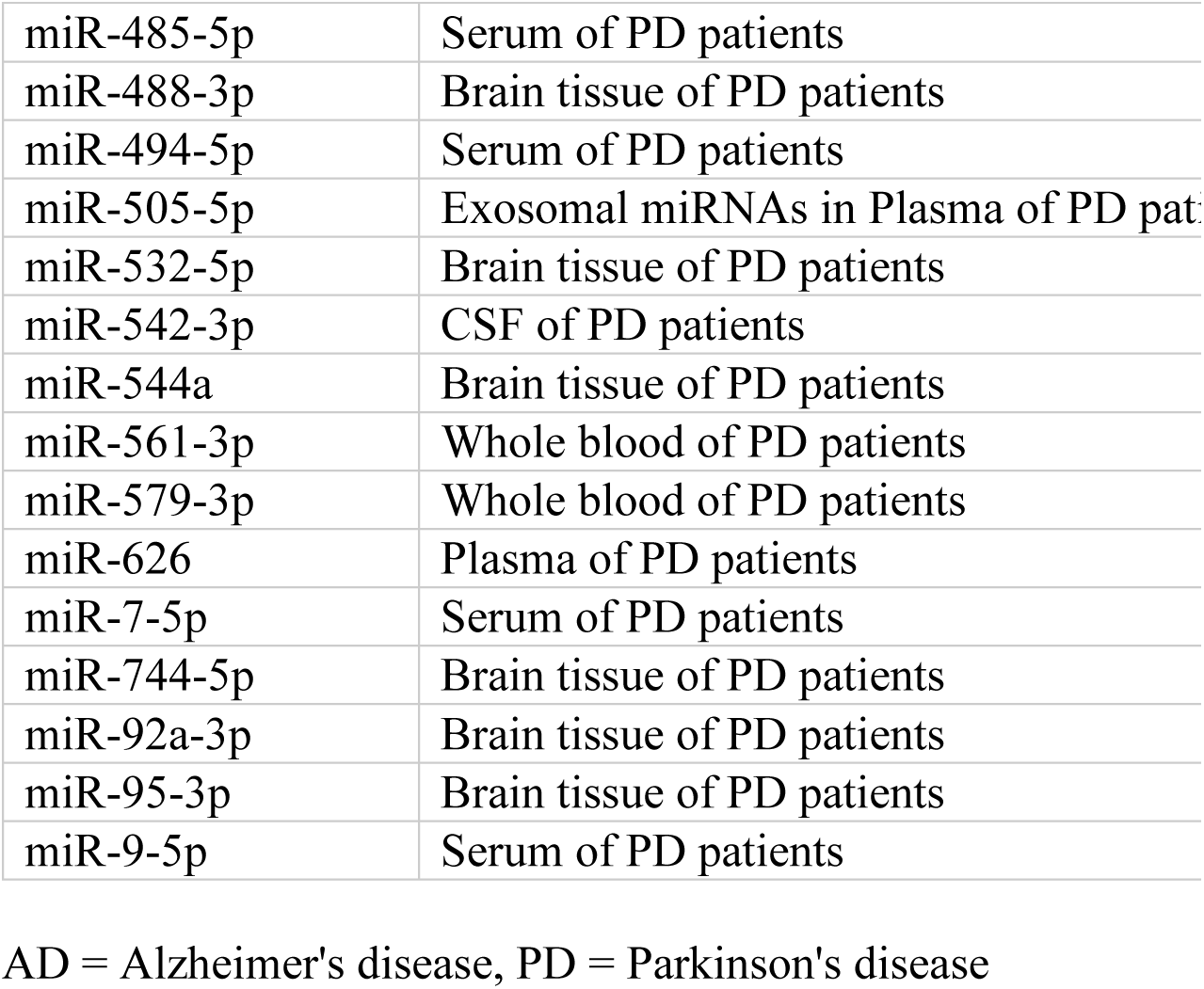
included miRNAs with literature references.

| **miRNA** | **Context** | **Reference** |
| --- | --- | --- |
| miR-16-5p | Striatal tissue of LRRK2 KO mice | [36] |
| miR-1-3p | Exosomal miRNAs in CSF of PD patients | [14] |
| miR-103a-3p | Striatal tissue of LRRK2 KO mice | [36] |
| miR-106a-5p | Brain tissue of PD patients | [30] |
| miR-107 | Algorithm based predicted interaction with LRRK2 | [37] |
|  | Plasma of AD and PD patients | [38] |
| miR-10a-5p | Exosomal miRNAs in CSF of PD patients | [14] |
| miR-122-5p | Striatal tissue of LRRK2 KO mice | [36] |
| miR-124-3p | Plasma of PD patients | [39] |
| miR-125b-5p | CSF of AD patients | [40] |
| miR-128-3p | Brain tissue of mouse models of PD | [41] |
| miR-129-5p | Algorithm based predicted interaction with LRRK2 | [37] |
| miR-132-3p | Brain tissue of PD patients | [30] |
| miR-133b | Plasma of PD patients | [42] |
| miR-135a-5p | Brain tissue of PD patients | [30] |
| miR-135b-5p | Brain tissue of PD patients | [43] |
| miR-136-5p | Algorithm based predicted interaction with LRRK2 | [37] |
| miR-137-3p | Plasma of PD patients | [39] |
| miR-141-5p | Algorithm based predicted interaction with LRRK2 | [37] |
| miR-144-5p | CSF of PD patients | [44] |
|  | Algorithm based predicted interaction with LRRK2 | [37] |
| miR-145-5p | Brain tissue of PD patients | [30] |
| miR-146a-5p | Serum of LRRK2 PD patients | [45] |
| miR-148a-3p | Brain tissue of PD patients | [30] |
| miR-153-3p | Exosomal miRNAs in CSF of PD patients | [14] |
| miR-155-5p | Serum of LRRK2 PD patients | [45] |
| miR-15b-5p | Serum of PD patients | [46] |
| miR-181a-5p | Serum of PD patients | [46] |
|  | Algorithm based predicted interaction with LRRK2 | [37] |
| miR-185-5p | Serum of PD patients | [46] |
|  | Algorithm based predicted interaction with LRRK2 | [37] |
| miR-190a-5p | Brain tissue of PD patients | [34] |
| miR-195-5p | Serum of PD patients | [46] |
| miR-198 | Brain tissue of PD patients | [43] |
| miR-199a-3p | Brain tissue of PD patients | [34] |
| miR-199b-5p | Brain tissue of PD patients | [47] |
| miR-19a-5p | Serum of LRRK2 PD patients | [26] |
|  | Algorithm based predicted interaction with LRRK2 | [37] |
| miR-19b-3p | Serum of PD patients | [26] |
|  | Algorithm based predicted interaction with LRRK2 | [37] |
|  | Exosomal miRNAs in CSF of PD patients | [14] |
| miR-200a-3p | CSF of PD patients | [44] |
|  | Algorithm based predicted interaction with LRRK2 | [37] |
| miR-200b-3p | Algorithm based predicted interaction with LRRK2 | [37] |
| miR-204-5p | Brain tissue of PD patients | [48] |
| miR-205-5p | CSF of PD patients | [49] |
|  | Algorithm based predicted interaction with LRRK2 | [37] |
| miR-214-5p | Serum of PD patients | [50] |
| miR-219a-2-3p | Brain tissue of PD patients | [48] |
| miR-221-3p | Plasma of PD patients | [42] |
| miR-222-3p | Plasma of PD patients | [51] |
| miR-223-3p | Brain tissue of PD patients | [30] |
| miR-24-3p | CSF of PD patients | [49] |
| miR-25-3p | Algorithm based predicted interaction with LRRK2 | [37] |
| miR-26a-5p | Brain tissue of PD patients | [30] |
| miR-27a-3p | CSF of PD patients | [52] |
| miR-28-5p | Brain tissue of PD patients | [30] |
| miR-29a-3p | Serum of LRRK2 PD patients | [26] |
| miR-29c-3p | Serum of LRRK2 PD patients | [26] |
| miR-301b-5p | Algorithm based predicted interaction with LRRK2 | [37] |
| miR-30b-5p | Serum of PD patients | [53] |
| miR-31-5p | Serum of PD advanced PD patients | [50] |
| miR-3143 | Whole blood of PD patients | [54] |
| miR-3200-3p | Brain tissue of PD patients | [48] |
| miR-328-5p | Algorithm based predicted interaction with LRRK2 | [37] |
| miR-331-5p | Exosomal miRNAs in Plasma of PD patients | [55] |
| miR-335-3p | Serum of LRRK2 PD patients | [45] |
| miR-335-5p | Serum of LRRK2 PD patients | [45] |
| miR-34b-5p | Brain tissue of PD patients | [56] |
| miR-34c-5p | Brain tissue of PD patients | [56] |
| miR-374a-5p | Brain tissue of PD patients | [30] |
| miR-382-5p | Brain tissue of PD patients | [48] |
| miR-384 | Algorithm based predicted interaction with LRRK2 | [37] |
| miR-409-3p | Exosomal miRNAs in CSF of PD patients | [14] |
| miR-410-5p | Algorithm based predicted interaction with LRRK2 | [37] |
| miR-421 | Brain tissue of PD patients | [48] |
| miR-423-5p | CSF of PD patients | [52] |
| miR-424-5p | Brain tissue of PD patients | [34] |
| miR-425-5p | Brain tissue of PD patients | [48] |
| miR-429 | Brain tissue of PD patients | [34] |
|  | Algorithm based predicted interaction with LRRK2 | [37] |
| miR-4421 | Brain tissue of PD patients | [48] |
| miR-451a | Brain tissue of PD patients | [34] |
| miR-454-5p | Algorithm based predicted interaction with LRRK2 | [37] |
| miR-4639-5p | Plasma of PD patients | [57] |
| miR-4671-3p | Whole blood of PD patients | [54] |
| miR-485-5p | Serum of PD patients | [58] |
| miR-488-3p | Brain tissue of PD patients | [47] |
| miR-494-5p | Serum of PD patients | [59] |
| miR-505-5p | Exosomal miRNAs in Plasma of PD patients | [55] |
| miR-532-5p | Brain tissue of PD patients | [30] |
| miR-542-3p | CSF of PD patients | [44] |
| miR-544a | Brain tissue of PD patients | [47] |
| miR-561-3p | Whole blood of PD patients | [54] |
| miR-579-3p | Whole blood of PD patients | [54] |
| miR-626 | Plasma of PD patients | [51] |
| miR-7-5p | Serum of PD patients | [60] |
| miR-744-5p | Brain tissue of PD patients | [30] |
| miR-92a-3p | Brain tissue of PD patients | [30] |
| miR-95-3p | Brain tissue of PD patients | [48] |
| miR-9-5p | Serum of PD patients | [61] |

AD = Alzheimer's disease, PD = Parkinson's disease

**Table S2:** Melting points of miRNAs detected in plasma. Only miRNAs which were selected for further analysis steps are displayed. Low standard deviations indicate constant amplification of the target.

| Target | Mean Melting Point [°C] | Standard deviation [°C] |
| --- | --- | --- |
| hsa-miR-103a-3p | 69.71 | 0.31 |
| hsa-miR-106a-5p | 71.71 | 0.34 |
| hsa-miR-107 | 69.73 | 0.34 |
| hsa-miR-10a-5p | 71.77 | 0.66 |
| hsa-miR-122-5p | 70.59 | 0.31 |
| hsa-miR-125b-5p | 71.37 | 0.38 |
| hsa-miR-128-3p | 69.75 | 0.11 |
| hsa-miR-132-3p | 71.20 | 0.16 |
| hsa-miR-133b | 71.54 | 0.21 |
| hsa-miR-135a-5p | 70.62 | 0.13 |
| hsa-miR-135b-5p | 69.36 | 0.32 |
| hsa-miR-136-5p | 70.30 | 0.13 |
| hsa-miR-1-3p | 70.65 | 0.13 |
| hsa-miR-144-5p | 71.39 | 0.18 |
| hsa-miR-145-5p | 69.09 | 0.32 |
| hsa-miR-146a-5p | 70.36 | 0.17 |
| hsa-miR-148a-3p | 69.78 | 0.27 |
| hsa-miR-153-3p | 69.92 | 0.22 |
| hsa-miR-155-5p | 69.63 | 0.19 |
| hsa-miR-15b-5p | 69.34 | 0.25 |
| hsa-miR-16-5p | 70.60 | 0.18 |
| hsa-miR-181a-5p | 70.26 | 0.17 |
| hsa-miR-185-5p | 70.65 | 0.16 |
| hsa-miR-190a-5p | 69.29 | 0.26 |
| hsa-miR-195-5p | 69.31 | 0.28 |
| hsa-miR-199a-3p | 69.94 | 0.13 |
| hsa-miR-19a-5p | 70.44 | 0.15 |
| hsa-miR-19b-3p | 69.88 | 0.15 |
| hsa-miR-200b-3p | 71.37 | 0.13 |
| hsa-miR-204-5p | 69.22 | 0.15 |
| hsa-miR-221-3p | 69.93 | 0.51 |
| hsa-miR-222-3p | 69.91 | 0.40 |
| hsa-miR-223-3p | 70.84 | 0.20 |
| hsa-miR-24-3p | 74.47 | 0.18 |
| hsa-miR-25-3p | 69.82 | 0.26 |
| hsa-miR-26a-5p | 70.88 | 0.12 |
| hsa-miR-27a-3p | 70.86 | 0.20 |
| hsa-miR-28-5p | 70.30 | 0.15 |
| hsa-miR-29a-3p | 70.19 | 0.23 |
| hsa-miR-29c-3p | 70.40 | 0.13 |
| hsa-miR-30b-5p | 69.20 | 0.10 |
| hsa-miR-31-5p | 69.22 | 1.82 |
| hsa-miR-335-3p | 70.89 | 0.13 |
| hsa-miR-335-5p | 71.43 | 0.25 |
| hsa-miR-374a-5p | 70.21 | 0.16 |
| hsa-miR-382-5p | 71.78 | 0.26 |
| hsa-miR-409-3p | 70.32 | 0.26 |
| hsa-miR-421 | 73.12 | 0.29 |
| hsa-miR-423-5p | 70.77 | 0.29 |
| hsa-miR-424-5p | 69.30 | 0.12 |
| hsa-miR-425-5p | 70.09 | 0.36 |
| hsa-miR-451a | 70.11 | 0.27 |
| hsa-miR-505-5p | 70.23 | 0.23 |
| hsa-miR-532-5p | 69.43 | 0.45 |
| hsa-miR-542-3p | 70.47 | 0.13 |
| hsa-miR-744-5p | 73.34 | 0.43 |
| hsa-miR-7-5p | 70.12 | 0.36 |
| hsa-miR-95-3p | 69.48 | 0.38 |
| hsa-miR-9-5p | 73.70 | 0.49 |

**Table S3:** Melting points of miRNAs detected in CSF. Only miRNAs which were selected for further analysis steps are displayed. Low standard deviations indicate constant amplification of the target.

| Target | Mean Melting Point [°C] | Standard deviation [°C] |
| --- | --- | --- |
| hsa-miR-16-5p | 70.84 | 0.47 |
| hsa-miR-125b-5p | 71.32 | 0.43 |
| hsa-miR-145-5p | 69.12 | 0.33 |
| hsa-miR-19b-3p | 69.75 | 0.27 |
| hsa-miR-204-5p | 69.25 | 0.15 |
| hsa-miR-223-3p | 70.66 | 0.38 |
| hsa-miR-24-3p | 74.19 | 1.05 |
| hsa-miR-26a-5p | 71.05 | 0.84 |
| hsa-miR-27a-3p | 70.83 | 0.29 |
| hsa-miR-29c-3p | 70.55 | 0.25 |
| hsa-miR-424-5p | 69.42 | 0.22 |

**Table S4:** Statistic details from plasma t-tests. Alpha was corrected to 0.0009 by dividing by the number of tests.

| **t-statistic** | **p-value** | **miRNA** |
| --- | --- | --- |
| -1.74 | 0.1019 | hsa-miR-1-3p |
| -2.10 | 0.0524 | hsa-miR-103a-3p |
| -2.72 | 0.0146 | hsa-miR-106a-5p |
| -2.50 | 0.0239 | hsa-miR-107 |
| -0.25 | 0.8073 | hsa-miR-10a-5p |
| 2.62 | 0.0218 | hsa-miR-122-5p |
| 0.72 | 0.4803 | hsa-miR-125b-5p |
| -3.06 | 0.0071 | hsa-miR-128-3p |
| -1.98 | 0.0658 | hsa-miR-132-3p |
| -1.39 | 0.1841 | hsa-miR-133b |
| -2.74 | 0.0161 | hsa-miR-135a-5p |
| -1.25 | 0.2296 | hsa-miR-135b-5p |
| -1.89 | 0.0764 | hsa-miR-136-5p |
| 1.37 | 0.1998 | hsa-miR-144-5p |
| -1.56 | 0.1384 | hsa-miR-145-5p |
| -1.90 | 0.0783 | hsa-miR-146a-5p |
| -3.31 | 0.0042 | hsa-miR-148a-3p |
| -3.58 | 0.0035 | hsa-miR-153-3p |
| -2.02 | 0.0597 | hsa-miR-155-5p |
| -1.57 | 0.1357 | hsa-miR-15b-5p |
| 0.08 | 0.9378 | hsa-miR-16-5p |
| -1.84 | 0.0851 | hsa-miR-181a-5p |
| -3.00 | 0.0081 | hsa-miR-185-5p |
| -2.23 | 0.0431 | hsa-miR-190a-5p |
| -2.60 | 0.0202 | hsa-miR-195-5p |
| -2.04 | 0.0588 | hsa-miR-199a-3p |
| -1.02 | 0.3222 | hsa-miR-19a-5p |
| -1.96 | 0.0734 | hsa-miR-19b-3p |
| -1.19 | 0.2502 | hsa-miR-200b-3p |
| -0.94 | 0.3590 | hsa-miR-204-5p |
| -1.37 | 0.1908 | hsa-miR-221-3p |
| -2.06 | 0.0574 | hsa-miR-222-3p |
| -2.10 | 0.0514 | hsa-miR-223-3p |
| -1.15 | 0.2666 | hsa-miR-24-3p |
| -0.82 | 0.4257 | hsa-miR-25-3p |
| -2.02 | 0.0658 | hsa-miR-26a-5p |
| -1.86 | 0.0822 | hsa-miR-27a-3p |
| -2.33 | 0.0358 | hsa-miR-28-5p |
| -1.73 | 0.1064 | hsa-miR-29a-3p |
| -4.21 | **0.0007\*** | hsa-miR-29c-3p |
| -2.07 | 0.0535 | hsa-miR-30b-5p |
| -1.55 | 0.1505 | hsa-miR-31-5p |
| -1.51 | 0.1489 | hsa-miR-335-3p |
| -2.24 | 0.0397 | hsa-miR-335-5p |
| -2.61 | 0.0186 | hsa-miR-374a-5p |
| -1.21 | 0.2421 | hsa-miR-382-5p |
| -1.27 | 0.2233 | hsa-miR-409-3p |
| -1.31 | 0.2068 | hsa-miR-421 |
| -2.35 | 0.0360 | hsa-miR-424-5p |
| -4.01 | 0.0010 | hsa-miR-425-5p |
| -0.62 | 0.5441 | hsa-miR-451a |
| -1.36 | 0.1945 | hsa-miR-505-5p |
| -2.08 | 0.0528 | hsa-miR-532-5p |
| 0.25 | 0.8022 | hsa-miR-542-3p |
| -1.26 | 0.2261 | hsa-miR-7-5p |
| -1.10 | 0.2890 | hsa-miR-744-5p |
| -2.75 | 0.0148 | hsa-miR-9-5p |
| 1.68 | 0.1106 | hsa-miR-95-3p |

**Table S5:** statistics of correlation matrix. Correlations calculated after Pearson. Only significant miRNA combinations (p < 0.05) are displayed.

| miRNA Plasma | miRNA CSF | R | R<sup>2</sup> | p-value |
| --- | --- | --- | --- | --- |
| miR-16-5p | miR-125-5p | 0.39 | 0.15 | 0.037 |
| miR-16-5p | miR-145-5p | 0.39 | 0.15 | 0.042 |
| miR-16-5p | miR-19b-3p | 0.40 | 0.16 | 0.034 |
| miR-16-5p | miR-223-3p | 0.43 | 0.19 | 0.021 |
| miR-16-5p | miR-27a-3p | 0.46 | 0.21 | 0.013 |
| miR-19b-3p | miR-125-5p | 0.38 | 0.15 | 0.041 |
| miR-19b-3p | miR-19b-3p | 0.44 | 0.20 | 0.018 |
| miR-19b-3p | miR-223-3p | 0.48 | 0.23 | 0.010 |
| miR-19b-3p | miR-27a-3p | 0.45 | 0.20 | 0.014 |
| miR-204-5p | miR-125b-5p | 0.37 | 0.14 | 0.043 |
| miR-204-5p | miR-19b-3p | 0.44 | 0.19 | 0.018 |
| miR-204-5p | miR-223-3p | 0.40 | 0.16 | 0.032 |
| miR-204-5p | miR-27a-3p | 0.48 | 0.23 | 0.007 |
| miR-204-5p | miR-29c-3p | 0.41 | 0.17 | 0.023 |
| miR-223-3p | miR-223-3p | 0.51 | 0.26 | 0.005 |
| miR-26a-5p | miR-223-3p | 0.49 | 0.24 | 0.007 |
| miR-26a-5p | miR-27a-3p | 0.36 | 0.13 | 0.032 |
| miR-27a-3p | miR-223-3p | 0.50 | 0.25 | 0.006 |
| miR-27a-3p | miR-27a-3p | 0.39 | 0.15 | 0.032 |
| miR-29c-3p | miR-27a-3p | 0.38 | 0.14 | 0.043 |

**Table S6:** t-test with Ct_multiplied_ data. p-value threshold was corrected for multiple testing by dividing by the number of tests and set to 0.0025.

| **miRNA Plasma** | **miRNA CSF** | **t-statistic** | **p-value** |
| --- | --- | --- | --- |
| miR-16-5p | miR-125-5p | 0.8 | 0.4483 |
| miR-16-5p | miR-145-5p | 1.1 | 0.3086 |
| miR-16-5p | miR-19b-3p | 2.0 | 0.0599 |
| miR-16-5p | miR-223-3p | 2.1 | 0.0500 |
| miR-16-5p | miR-27a-3p | 1.4 | 0.1898 |
| miR-19b-3p | miR-125-5p | 1.5 | 0.1602 |
| miR-19b-3p | miR-19b-3p | 2.8 | 0.0132 |
| miR-19b-3p | miR-223-3p | 2.8 | 0.0119 |
| miR-19b-3p | miR-27a-3p | 2.1 | 0.0544 |
| miR-204-5p | miR-125b-5p | 1.0 | 0.3343 |
| miR-204-5p | miR-19b-3p | 2.1 | 0.0517 |
| miR-204-5p | miR-223-3p | 2.3 | 0.0324 |
| miR-204-5p | miR-27a-3p | 1.5 | 0.1457 |
| miR-204-5p | miR-29c-3p | 1.3 | 0.2131 |
| miR-223-3p | miR-223-3p | 2.9 | 0.0113 |
| miR-26a-5p | miR-223-3p | 2.6 | 0.0223 |
| miR-26a-5p | miR-27a-3p | 2.2 | 0.0467 |
| miR-27a-3p | miR-223-3p | 2.6 | 0.0204 |
| miR-27a-3p | miR-27a-3p | 2.1 | 0.0532 |
| miR-29c-3p | miR-27a-3p | 4.2 | **0.0007\*** |

## References

[1] Zimprich A, Biskup S, Leitner P, Lichtner P, Farrer M, Lincoln S, Kachergus J, Hulihan M, Uitti RJ, Calne DB, Stoessl AJ, Pfeiffer RF, Patenge N, Carbajal IC, Vieregge P, Asmus F, Müller-Myhsok B, Dickson DW, Meitinger T, Strom TM, Wszolek ZK, Gasser T (2004) Mutations in LRRK2 Cause Autosomal-Dominant Parkinsonism with Pleomorphic Pathology. Neuron 44, 601–607.

[2] Paisán-Ruíz C, Jain S, Evans EW, Gilks WP, Simón J, van der Brug M, de Munain AL, Aparicio S, Gil AM, Khan N, Johnson J, Martinez JR, Nicholl D, Carrera IM, Peňa AS, de Silva R, Lees A, Martí-Massó JF, Pérez-Tur J, Wood NW, Singleton AB (2004) Cloning of the Gene Containing Mutations that Cause PARK8-Linked Parkinson’s Disease. Neuron 44, 595–600.

[3] Correia Guedes L, Ferreira JJ, Rosa MM, Coelho M, Bonifati V, Sampaio C (2010) Worldwide frequency of G2019S LRRK2 mutation in Parkinson’s disease: A systematic review. Parkinsonism Relat Disord 16, 237–242.

[4] Uhlén M, Fagerberg L, Hallström BM, Lindskog C, Oksvold P, Mardinoglu A, Sivertsson Å, Kampf C, Sjöstedt E, Asplund A, Olsson I, Edlund K, Lundberg E, Navani S, Szigyarto CA-K, Odeberg J, Djureinovic D, Takanen JO, Hober S, Alm T, Edqvist P-H, Berling H, Tegel H, Mulder J, Rockberg J, Nilsson P, Schwenk JM, Hamsten M, von Feilitzen K, Forsberg M, Persson L, Johansson F, Zwahlen M, von Heijne G, Nielsen J, Pontén F (2015) Tissue-based map of the human proteome. Science (80-) 347,.

[5] The Human Protein Atlas.

[6] Jaleel M, Nichols RJ, Deak M, Campbell DG, Gillardon F, Knebel A, Alessi DR (2007) LRRK2 phosphorylates moesin at threonine-558: characterization of how Parkinson’s disease mutants affect kinase activity. Biochem J 405, 307–317.

[7] Simón-Sánchez J, Schulte C, Bras JM, Sharma M, Gibbs JR, Berg D, Paisan-Ruiz C, Lichtner P, Scholz SW, Hernandez DG, Krüger R, Federoff M, Klein C, Goate A, Perlmutter J, Bonin M, Nalls MA, Illig T, Gieger C, Houlden H, Steffens M, Okun MS, Racette BA, Cookson MR, Foote KD, Fernandez HH, Traynor BJ, Schreiber S, Arepalli S, Zonozi R, Gwinn K, van der Brug M, Lopez G, Chanock SJ, Schatzkin A, Park Y, Hollenbeck A, Gao J, Huang X, Wood NW, Lorenz D, Deuschl G, Chen H, Riess O, Hardy JA, Singleton AB, Gasser T (2009) Genome-wide association study reveals genetic risk underlying Parkinson’s disease. Nat Genet 41, 1308–1312.

[8] Schiesling C, Kieper N, Seidel K, Krüger R (2008) Review: Familial Parkinson’s disease – genetics, clinical phenotype and neuropathology in relation to the common sporadic form of the disease. Neuropathol Appl Neurobiol 34, 255–271.

[9] Kalia L V, Lang AE (2015) Parkinson’s disease. *Lancet (London*, England) 386, 896–912.

[10] Azeggagh S, Berwick DC (2022) The development of inhibitors of leucine-rich repeat kinase 2 (LRRK2) as a therapeutic strategy for Parkinson’s disease: the current state of play. Br J Pharmacol 179, 1478–1495.

[11] Jennings D, Huntwork-Rodriguez S, Henry AG, Sasaki JC, Meisner R, Diaz D, Solanoy H, Wang X, Negrou E, Bondar V V., Ghosh R, Maloney MT, Propson NE, Zhu Y, Maciuca RD, Harris L, Kay A, LeWitt P, King TA, Kern D, Ellenbogen A, Goodman I, Siderowf A, Aldred J, Omidvar O, Masoud ST, Davis SS, Arguello A, Estrada AA, de Vicente J, Sweeney ZK, Astarita G, Borin MT, Wong BK, Wong H, Nguyen H, Scearce-Levie K, Ho C, Troyer MD (2022) Preclinical and clinical evaluation of the LRRK2 inhibitor DNL201 for Parkinson’s disease. Sci Transl Med 14,.

[12] Bartel DP (2004) MicroRNAs: genomics, biogenesis, mechanism, and function. Cell 116, 281–297.

[13] Burgos K, Malenica I, Metpally R, Courtright A, Rakela B, Beach T, Shill H, Adler C, Sabbagh M, Villa S, Tembe W, Craig D, Van Keuren-Jensen K (2014) Profiles of Extracellular miRNA in Cerebrospinal Fluid and Serum from Patients with Alzheimer’s and Parkinson’s Diseases Correlate with Disease Status and Features of Pathology. PLoS One 9, e94839.

[14] Gui Y, Liu H, Zhang L, Lv W, Hu X (2015) Altered microRNA profiles in cerebrospinal fluid exosome in Parkinson disease and Alzheimer disease. Oncotarget 6, 37043–37053.

[15] Gehrke S, Imai Y, Sokol N, Lu B (2010) Pathogenic LRRK2 negatively regulates microRNA-mediated translational repression. Nature 466, 637–641.

[16] Ramakers C, Ruijter JM, Deprez RHL, Moorman AF. (2003) Assumption-free analysis of quantitative real-time polymerase chain reaction (PCR) data. Neurosci Lett 339, 62–66.

[17] R Core Team (2022) A language and environment for statistical computing.

[18] Livak KJ, Schmittgen TD (2001) Analysis of Relative Gene Expression Data Using Real-Time Quantitative PCR and the 2−ΔΔCT Method. Methods 25, 402– 408.

[19] Kolde R (2019) pheatmap: Pretty Heatmaps.

[20] Kuhn M (2008) Building Predictive Models in R Using the caret Package. J Stat Softw 28,.

[21] Kohavi R (2001) A Study of Cross-Validation and Bootstrap for Accuracy Estimation and Model Selection. 14,.

[22] Liaw A, Wiener M (2002) Classification and Regression by randomForest. R News 2, 18–22.

[23] Chen Y, Wang X (2020) miRDB: an online database for prediction of functional microRNA targets. Nucleic Acids Res 48, D127–D131.

[24] Wu T, Hu E, Xu S, Chen M, Guo P, Dai Z, Feng T, Zhou L, Tang W, Zhan L, Fu X, Liu S, Bo X, Yu G (2021) clusterProfiler 4.0: A universal enrichment tool for interpreting omics data. Innov 2, 100141.

[25] Wang R, Yang Y, Wang H, He Y, Li C (2020) MiR-29c protects against inflammation and apoptosis in Parkinson’s disease model in vivo and in vitro by targeting SP1. Clin Exp Pharmacol Physiol 47, 372–382.

[26] Botta-Orfila T, Morató X, Compta Y, Lozano JJ, Falgàs N, Valldeoriola F, Pont-Sunyer C, Vilas D, Mengual L, Fernández M, Molinuevo JL, Antonell A, Martí MJ, Fernández-Santiago R, Ezquerra M (2014) Identification of blood serum micro-RNAs associated with idiopathic and LRRK2 Parkinson’s disease. J Neurosci Res 92, 1071–1077.

[27] Bai X, Tang Y, Yu M, Wu L, Liu F, Ni J, Wang Z, Wang J, Fei J, Wang W, Huang F, Wang J (2017) Downregulation of blood serum microRNA 29 family in patients with Parkinson’s disease. Sci Rep 7, 5411.

[28] Ozdilek B, Demircan B (2021) Serum microRNA expression levels in Turkish patients with Parkinson’s disease. Int J Neurosci 131, 1181–1189.

[29] Soto M, Fernández M, Bravo P, Lahoz S, Garrido A, Sánchez-Rodríguez A, Rivera-Sánchez M, Sierra M, Melón P, Roig-García A, Naito A, Casey B, Camps J, Tolosa E, Martí M-J, Infante J, Ezquerra M, Fernández-Santiago R (2023) Differential serum microRNAs in premotor LRRK2 G2019S carriers from Parkinson’s disease. npj Park Dis 9, 15.

[30] Briggs CE, Wang Y, Kong B, Woo T-UW, Iyer LK, Sonntag KC (2015) Midbrain dopamine neurons in Parkinson׳s disease exhibit a dysregulated miRNA and target-gene network. Brain Res 1618, 111–121.

[31] Mancuso R, Agostini S, Hernis A, Zanzottera M, Bianchi A, Clerici M (2019) Circulatory miR-223-3p Discriminates Between Parkinson’s and Alzheimer’s Patients. Sci Rep 9, 9393.

[32] Bauernfeind F, Rieger A, Schildberg FA, Knolle PA, Schmid-Burgk JL, Hornung V (2012) NLRP3 Inflammasome Activity Is Negatively Controlled by miR-223. J Immunol 189, 4175–4181.

[33] Moehle MS, Webber PJ, Tse T, Sukar N, Standaert DG, DeSilva TM, Cowell RM, West AB (2012) LRRK2 Inhibition Attenuates Microglial Inflammatory Responses. J Neurosci 32, 1602–1611.

[34] Thomas RR, Keeney PM, Bennett JP (2012) Impaired Complex-I Mitochondrial Biogenesis in Parkinson Disease Frontal Cortex. J Parkinsons Dis 2, 67–76.

[35] Kanao T, Venderova K, Park DS, Unterman T, Lu B, Imai Y (2010) Activation of FoxO by LRRK2 induces expression of proapoptotic proteins and alters survival of postmitotic dopaminergic neuron in Drosophila. Hum Mol Genet 19, 3747–3758.

[36] Dorval V, Mandemakers W, Jolivette F, Coudert L, Mazroui R, Strooper B, Hébert SS (2014) Gene and MicroRNA transcriptome analysis of Parkinson’s related LRRK2 mouse models. PLoS One 9, e85510.

[37] Heman-Ackah SM, Hallegger M, Rao MS, Wood MJA (2013) RISC in PD: the impact of microRNAs in Parkinson’s disease cellular and molecular pathogenesis. Front Mol Neurosci 6,.

[38] Wang J, Chen C, Zhang Y (2020) An investigation of microRNA-103 and microRNA-107 as potential blood-based biomarkers for disease risk and progression of Alzheimer’s disease. J Clin Lab Anal 34,.

[39] Li N, Pan X, Zhang J, Ma A, Yang S, Ma J, Xie A (2017) Plasma levels of miR-137 and miR-124 are associated with Parkinson’s disease but not with Parkinson’s disease with depression. Neurol Sci 38, 761–767.

[40] Alexandrov PN, Dua P, Hill JM, Bhattacharjee S, Zhao Y, Lukiw WJ (2012) microRNA (miRNA) speciation in Alzheimer’s disease (AD) cerebrospinal fluid (CSF) and extracellular fluid (ECF). Int J Biochem Mol Biol 3, 365–73.

[41] Zhou L, Yang L, Li Y, Mei R, Yu H, Gong Y, Du M, Wang F (2018) MicroRNA-128 Protects Dopamine Neurons from Apoptosis and Upregulates the Expression of Excitatory Amino Acid Transporter 4 in Parkinson’s Disease by Binding to AXIN1. Cell Physiol Biochem 51, 2275–2289.

[42] Chen Q, Deng N, Lu K, Liao Q, Long X, Gou D, Bi F, Zhou J (2021) Elevated plasma miR-133b and miR-221-3p as biomarkers for early Parkinson’s disease. Sci Rep 11, 15268.

[43] Cardo LF, Coto E, Ribacoba R, Menéndez M, Moris G, Suárez E, Alvarez V (2014) MiRNA Profile in the Substantia Nigra of Parkinson’s Disease and Healthy Subjects. J Mol Neurosci 54, 830–836.

[44] Mo M, Xiao Y, Huang S, Cen L, Chen X, Zhang L, Luo Q, Li S, Yang X, Lin X, Xu P (2017) MicroRNA expressing profiles in A53T mutant alpha-synuclein transgenic mice and Parkinsonian. Oncotarget 8, 15–28.

[45] Oliveira SR, Dionísio PA, Correia Guedes L, Gonçalves N, Coelho M, Rosa MM, Amaral JD, Ferreira JJ, Rodrigues CMP (2020) Circulating Inflammatory miRNAs Associated with Parkinson’s Disease Pathophysiology. Biomolecules 10,.

[46] Ding H, Huang Z, Chen M, Wang C, Chen X, Chen J, Zhang J (2016) Identification of a panel of five serum miRNAs as a biomarker for Parkinson’s disease. Parkinsonism Relat Disord 22, 68–73.

[47] Tatura R, Kraus T, Giese A, Arzberger T, Buchholz M, Höglinger G, Müller U (2016) Parkinson’s disease: SNCA-, PARK2-, and LRRK2-targeting microRNAs elevated in cingulate gyrus. Parkinsonism Relat Disord 33, 115–121.

[48] Nair VD, Ge Y (2016) Alterations of miRNAs reveal a dysregulated molecular regulatory network in Parkinson’s disease striatum. Neurosci Lett 629, 99–104.

[49] Marques TM, Kuiperij HB, Bruinsma IB, van Rumund A, Aerts MB, Esselink RAJ, Bloem BR, Verbeek MM (2017) MicroRNAs in Cerebrospinal Fluid as Potential Biomarkers for Parkinson’s Disease and Multiple System Atrophy. Mol Neurobiol 54, 7736–7745.

[50] Li L, Ren J, Pan C, Li Y, Xu J, Dong H, Chen Y, Liu W (2021) Serum miR-214 Serves as a Biomarker for Prodromal Parkinson’s Disease. Front Aging Neurosci 13,.

[51] Khoo SK, Petillo D, Kang UJ, Resau JH, Berryhill B, Linder J, Forsgren L, Neuman LA, Tan AC (2012) Plasma-Based Circulating MicroRNA Biomarkers for Parkinson’s Disease. J Parkinsons Dis 2, 321–331.

[52] dos Santos MCT, Barreto-Sanz MA, Correia BRS, Bell R, Widnall C, Perez LT, Berteau C, Schulte C, Scheller D, Berg D, Maetzler W, Galante PAF, Nogueira da Costa A (2018) miRNA-based signatures in cerebrospinal fluid as potential diagnostic tools for early stage Parkinson’s disease. Oncotarget 9, 17455–17465.

[53] Han K (2017) The diagnostic efficacy of serum miR-103a, miR-30b, miR-29a relative expression levels for Parkinson’s disease. Shadong Med J 57, 72–74.

[54] Yılmaz ŞG, Geyik S, Neyal AM, Soko ND, Bozkurt H, Dandara C (2016) Hypothesis: Do miRNAs Targeting the Leucine-Rich Repeat Kinase 2 Gene (*LRRK2*) Influence Parkinson’s Disease Susceptibility? Omi A J Integr Biol 20, 224–228.

[55] Yao Y-F, Qu M-W, Li G-C, Zhang F-B, Rui H-C (2018) Circulating exosomal miRNAs as diagnostic biomarkers in Parkinson’s disease. Eur Rev Med Pharmacol Sci 22, 5278–5283.

[56] Miñones-Moyano E, Porta S, Escaramís G, Rabionet R, Iraola S, Kagerbauer B, Espinosa-Parrilla Y, Ferrer I, Estivill X, Martí E (2011) MicroRNA profiling of Parkinson’s disease brains identifies early downregulation of miR-34b/c which modulate mitochondrial function. Hum Mol Genet 20, 3067–3078.

[57] Chen Y, Gao C, Sun Q, Pan H, Huang P, Ding J, Chen S (2017) MicroRNA-4639 Is a Regulator of DJ-1 Expression and a Potential Early Diagnostic Marker for Parkinson’s Disease. Front Aging Neurosci 9,.

[58] Lin X, Wang R, Li R, Tao T, Zhang D, Qi Y (2022) Diagnostic Performance of miR-485-3p in Patients with Parkinson’s Disease and its Relationship with Neuroinflammation. NeuroMolecular Med 24, 195–201.

[59] Li J, Yin Y, Wang R, Zhang P, Zhao J, Liu R, Chen Y (2021) Expressions and clinical significance of miR-124 and miR-494 in elderly patients with Parkinson disease. Chinese J Behav Med Brain Sci 12, 294–298.

[60] Zhou Y, Lu M, Du R-H, Qiao C, Jiang C-Y, Zhang K-Z, Ding J-H, Hu G (2016) MicroRNA-7 targets Nod-like receptor protein 3 inflammasome to modulate neuroinflammation in the pathogenesis of Parkinson’s disease. Mol Neurodegener 11, 28.

[61] Wu L, Zhao W, Kong F, Yang F, Zheng J (2020) SERUM MIR-9A AND MIR-133B, DIAGNOSTIC MARKERS FOR PARKINSON’S DISEASE, ARE UPREGULATED AFTER LEVODOPA TREATMENT. Acta Medica Mediterr 36,.

